# *HDAC9* structural variants disrupting *TWIST1* transcriptional regulation lead to craniofacial and limb malformations

**DOI:** 10.1101/2021.08.10.455254

**Authors:** Naama Hirsch, Idit Dahan, Eva D’haene, Matan Avni, Sarah Vergult, Marta Vidal-García, Pamela Magini, Claudio Graziano, Giulia Severi, Elena Bonora, Anna Maria Nardone, Francesco Brancati, Alberto Fernández-Jaén, Olson J. Rory, Benedikt Hallgrimsson, Ramon Y. Birnbaum

**Author notes:** Corresponding author: Ramon Y. Birnbaum.

## Abstract

Structural variants (SVs) can affect protein-coding sequences as well as gene regulatory elements. However, SVs disrupting protein-coding sequences that also function as cis-regulatory elements remain largely uncharacterized. Here, we show that craniosynostosis patients with SVs containing the Histone deacetylase 9 (*HDAC9*) protein-coding sequence are associated with disruption of *TWIST1* regulatory elements that reside within *HDAC9* sequence. Based on SVs within the *HDAC9-TWIST1* locus, we defined the 3’ HDAC9 sequence (~500Kb) as a critical *TWIST1* regulatory region, encompassing craniofacial *TWIST1* enhancers and CTCF sites. Deletions of either *Twist1* enhancers (**eTw5-7^Δ/Δ^**) or Ctcf site (**Ctcf^Δ/Δ^**) within the Hdac9 protein-coding sequence in mice led to decreased *Twist1* expression and altered anterior\posterior limb expression patterns of Shh pathway genes. This decreased Twist1 expression results in a smaller sized and asymmetric skull and polydactyly that resembles *Twist1*^+/-^ mouse phenotype. Chromatin conformation analysis revealed that *the Twist1* promoter region interacts with *Hdac9* sequences that encompass *Twist1* enhancers and a Ctcf site and that interactions depended on the presence of both regulatory regions. Finally, a large inversion of the entire *Hdac9* sequence (***Hdac9*^INV/+^**) in mice that does not disrupt *Hdac9* expression but repositions *Twist1* regulatory elements showed decreased *Twist1* expression and led to a craniosynostosis-like phenotype and polydactyly. Thus, our study elucidated essential components of *TWIST1* transcriptional machinery that reside within the *HDAC9* sequence, suggesting that SVs, encompassing protein-coding sequence, such as *HDAC9*, could lead to a phenotype that is not attributed to its protein function but rather to a disruption of the transcriptional regulation of a nearby gene, such as *TWIST1*.

## Introduction

Structural variants (SVs) involve at least 50 nucleotides, rearranging large segments of DNA and often have profound consequences in evolution and human disease^1,2^. Given their size and abundance, SVs represent an important mutational force that shapes genome function and contributes to germline and somatic diseases. A recent study analyzing ~15,000 genomes discovered that SVs are responsible for ~25% of all rare protein-truncating events per genome, indicating that SVs have a major effect on protein sequences^1^. Moreover, an underscored modest selection was found against noncoding SVs in cis-regulatory elements that control spatiotemporal gene expression. The profound effect of SVs is also attributable to the numerous mechanisms by which they disrupt protein-coding genes and cis-regulatory architecture^3^. SVs can alter the copy number of regulatory elements or 3D genome structure by disrupting higher-order chromatin organization such as topologically associating domains (TADs)^4^. As a result of these position effects, SVs can also influence the expression of genes distant from the SV breakpoints, causing disease. However, the effect of SVs on gene regulatory mechanisms remains poorly understood.

We have previously demonstrated that protein-coding DNA sequences could also function as enhancers of nearby genes, indicating a dual function of DNA sequences ^5–9^. We have studied the *TWIST1-HDAC9* locus as an example of this dual function of DNA sequences. TWIST1 is a transcription factor (TF) that plays a critical role in mesodermal development^10^. TWIST1 regulates the expression of various other TFs and signaling pathways in the developing craniofacial and limb tissues and *TWIST1* haploinsufficiency is associated with a range of craniofacial and limb malformations^11^. *TWIST1* protein-coding mutations, including deletions, are associated with Saethre-Chotzen syndrome (OMIM #101400), an autosomal dominant craniosynostosis disorder (pre-mature closer of the sutures) also associated with distal limb malformations^12,13^. Previous studies have shown that *Twist1* homozygous null mice are lethal, but *Twist1* heterozygous mice exhibit a craniosynostosis-like phenotype along with polydactyly^11,14^. These findings indicate the existence of a *Twist1* dosage threshold that is likely regulated by tissue-specific enhancers that is essential for correct craniofacial and limb formation. Recently, we identified tissue-specific enhancers located in the *HDAC9-TWIST1* locus that recapitulate *Twist1* expression^8^ and comprising a spatiotemporal regulatory network of *Twist1* transcription in the developing limbs/fins^8^.

Histone deacetylase 9 (HDAC9) encodes an enzyme that modifies the N-terminal tail of histones and leads to compact chromatin structure and reduced gene expression^15^. The Hdac9 protein is highly expressed in mouse adult brain and heart tissues^16–19^. In the brain, Hdac9 expression is limited to post-mitotic neurons and highly expressed in the hippocampus and cerebral cortex^18^. Copy number variants of HDAC9 are associated with schizophrenia in humans and with neuropathological changes in the hippocampus and cerebral cortex in mice^18^. Hdac9 also inhibits skeletal myogenesis and is involved in heart development^20^. While *Hdac9*-null mice are fertile and have a normal life span, they develop cardiac hypertrophy with age and in response to pressure overload^17^. Interestingly, a polydactyly phenotype has also been discovered in *Hdac9* null mice^21^. However, *Hdac9* is not expressed in the developing limb, suggesting that the polydactyly phenotype in these mice is independent of the Hdac9 protein function^8^. Instead, the polydactyly phenotype could result from a disruption of *Twist1* regulatory elements residing in the *Hdac9* sequence, leading to haploinsufficiency of *Twist1*.

Here, we study the effects of disruption of protein-coding sequences located in a critical regulatory region for a nearby gene. We identified and characterized craniofacial enhancers and bound CTCF regions in the *HDAC9* sequence. These regulatory elements are part of a critical TWIST1 regulatory region that is disrupted by *HDAC9*-encompassing SVs in patients with craniosynostosis. We then modeled these *HDAC9* SVs in mice, which resulted in craniosynostosis-like phenotype and polydactyly. The phenotype correlated with reduced frequency of chromatin interactions between *Twist1* regulatory elements and with reduced *Twist1* expression, leading to Twist1 haploinsufficiency. Thus, *HDAC9*-encompassing SVs cause craniosynostosis through the disruption of a critical TWIST1 regulatory region that resides within the HDAC9 sequence.

## Results

### *TWIST1* craniofacial enhancers reside within the *HDAC9* sequence

We previously described *TWIST1* limb enhancers during embryonic development and characterized their function and activity^8^. Nevertheless, the regulatory mechanisms controlling *TWIST1* expression in craniofacial tissues are barely known. To explore the regulatory elements of *TWIST1* during craniofacial development and skull formation, we focused on the *TWIST1-HDAC9* locus (hg19: chr7:18,050,988–19,741,484) and analyzed chromatin immunoprecipitation sequencing (ChIP-seq) of multiple histone modifications (H3K4me1, H3K4me2, H3K4me3, H3K36me3, H3K27ac, H3K27me3) from early human embryonic craniofacial tissues (stages CS13-CS14, corresponding to mouse embryonic day10.5-11.5)^22^. Analyzing these datasets, we identified 25 sequences as active enhancer candidates. We further analyzed ATAC-seq and histone modification ChIP-seq of mouse E10.5 craniofacial tissues (maxilla, mandibula, pharyngeal arch 2, and frontal nasal plate)^23^, searching for sequences marked as active enhancers in mouse craniofacial developmental tissues. We defined sequences as enhancer candidates if they were necessarily marked in CS13-CS14 human data as well as in at least one of the mouse datasets. These 15 sequences, corresponding to 5 intergenic, 6 *HDAC9* intronic, and 4 *HDAC9* exonic sequences, might regulate *TWIST1* transcription during craniofacial development **(Fig. S1, Table S1)**. Next, we used unique molecular identifier (UMI) circularized chromosome conformation capture sequencing (UMI-4C) on mouse E11.5 branchial arches 1-2 (BA) to determine chromatin interactions between the *Twist1* promoter region and candidate enhancers. Using the *Twist1* promoter as a viewpoint, we found that *Twist1* frequently interacts with several regions encompassing enhancer candidates (eTw5, eTw6, eTw18, and eTw19) and Ctcf sites **(Fig. 1A-B)**, supporting that these enhancer candidates regulate *Twist1* expression. Then, we tested these candidates for functional activity using the transgenic zebrafish enhancer assay. Of the 15 sequences, we previously reported on four sequences (eTw2, eTw5, eTw6, eTw11) as craniofacial and/or limb enhancers **(Table S2)**^8^. The zebrafish enhancer assay showed that eTw2 and eTw6 drove specific GFP expression in branchial arches 1-2 (BA1-2), which are homologous to mammalian mandibular arch, maxillar arch, and hyoid. eTw5 displayed a broad activity pattern in branchial arches and drove GFP in branchial arches 1-7 (BA1-7). eTw11 drove GFP expression in the posterior part of the branchial arches 3-7 (BA3-7) **(Fig. 1C)**. Two additional craniofacial enhancer candidates, eTw18 and eTw19, were shown to have functional activity in the craniofacial tissues of zebrafish embryos **(Fig. 1C)**. eTw18 drove GFP expression in the branchial arches 1-7 with similarity to eTw5 and eTw19 drove GFP expression in branchial arches 1-7 and also in the front nasal and maxillary prominences **(Fig 1C; Fig. S2)**. Thus, each enhancer has a discrete activity pattern, and together comprise a spatiotemporal regulatory network that likely controls Twist1 expression in the developing craniofacial tissues **(Table S2)**.

**Figure 1:**
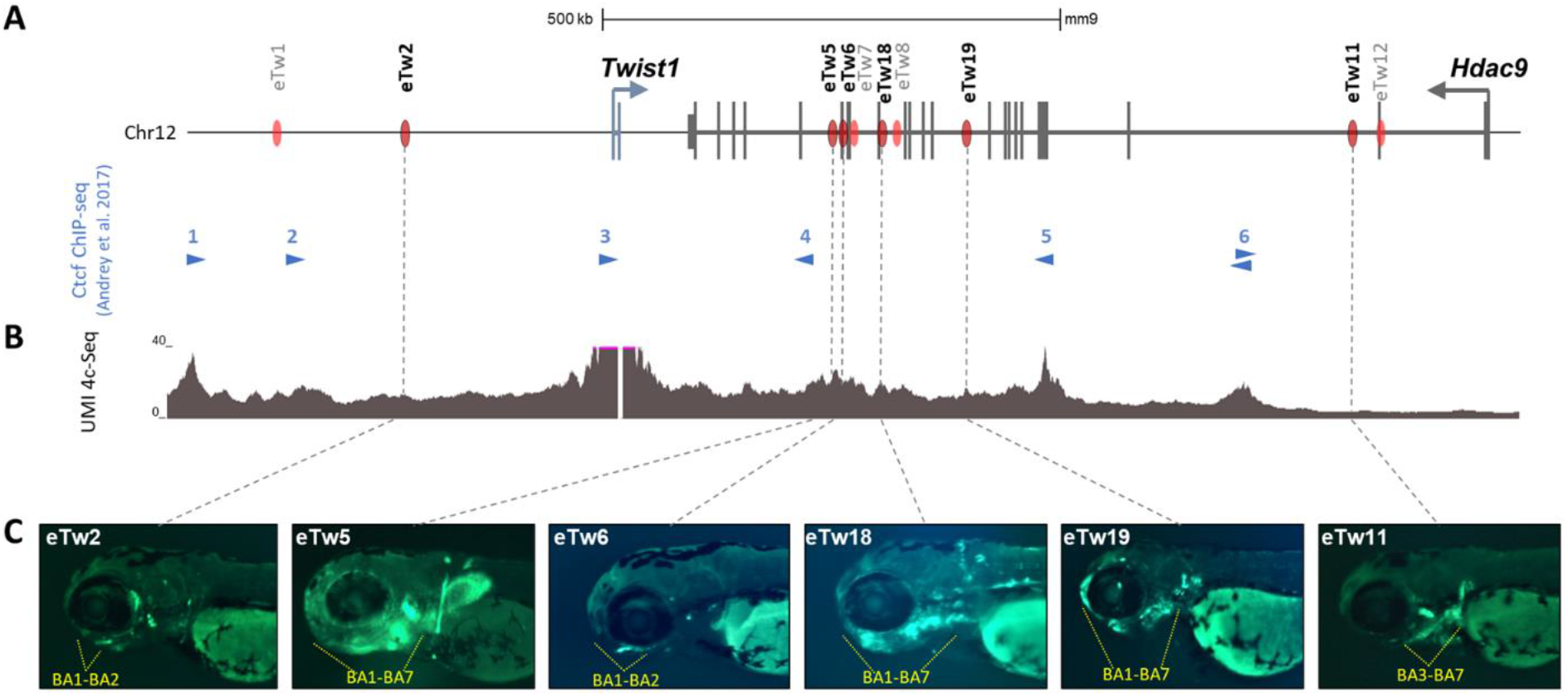
*Twist1* craniofacial enhancers in the *Hdac9-Twist1* locus. **(A)** A scheme of the *Hdac9-Twist1* locus. Black lines represent exons. Red ovals represent positive enhancer sequences in zebrafish enhancer assay. Blue arrowheads represent the directionality of Ctcf bound sites in E11.5 limb buds^24^. **(B)** UMI-4C interaction profile (based on 2 biological replicates) using the *Twist1* promoter as a viewpoint in the branchial arches of mouse E11.5 embryos. **(C)** Activity pattern of Twist1 craniofacial enhancer in zebrafish. As zebrafish has 7 branchial arches (BA1-7), yellow dash lines show the GFP expression patterns in the branchial arches of zebrafish embryos at 72hpf.

### Structural variants compromise the HDAC9 coding sequence in patients with craniofacial malformations

To demonstrate that disruption of protein-coding sequences can also affect regulatory elements of nearby genes, we collected patients with SVs in *HDAC9*. Through the international Matchmaker Exchange initiative^25^, we found craniofacial malformation patients with deletions containing *HDAC9* coding sequences that also function as regulatory elements of the neighboring *TWIST1* gene **(Table S3, Fig. 2)**. Two craniosynostosis patients were reported with an *HDAC9* deletion (P1) and a translocation with an intergenic breakpoint between *HDAC9* and *TWIST1* (*P4*), respectively^26,27^. We found two additional craniosynostosis patients with *HDAC9* deletions, in which the *TWIST1* protein-coding sequence is not disrupted (P2-P3, **Fig. 2)**. Conversely, other *HDAC9* deletions overlapping the 5’ region of *HDAC9* are associated with neuronal disorders, such as schizophrenia^18^, without reported craniofacial phenotype (P5-P7, **Fig. 2**). In addition, we identified *HDAC9* single nucleotide variants (SNVs), including a splice site, frameshift, and missense variant in patients with global developmental delay, thin corpus callosum, and seizures, but without craniosynostosis **(P8-10, Table S3)**. The enrichment for loss of function variants in our cohort is consistent with the constraint data from the Exome Aggregation Consortium (ExAC) database, suggesting that HDAC9 is extremely intolerant to loss-of-function variations (probability of being loss-of-function intolerant (pLI) = 1)^28^. Therefore, we concluded that SVs affecting the 3’ sequence of *HDAC9*, in the region delineated by two convergent CTCF sites (e.g. 3 and 5) and containing *TWIST1* enhancers, are likely associated with craniofacial malformations, suggesting that the 3’ terminal part of *HDAC9* is critical for *TWIST1* transcriptional regulation **(Fig. 2)**.

**Figure 2.**
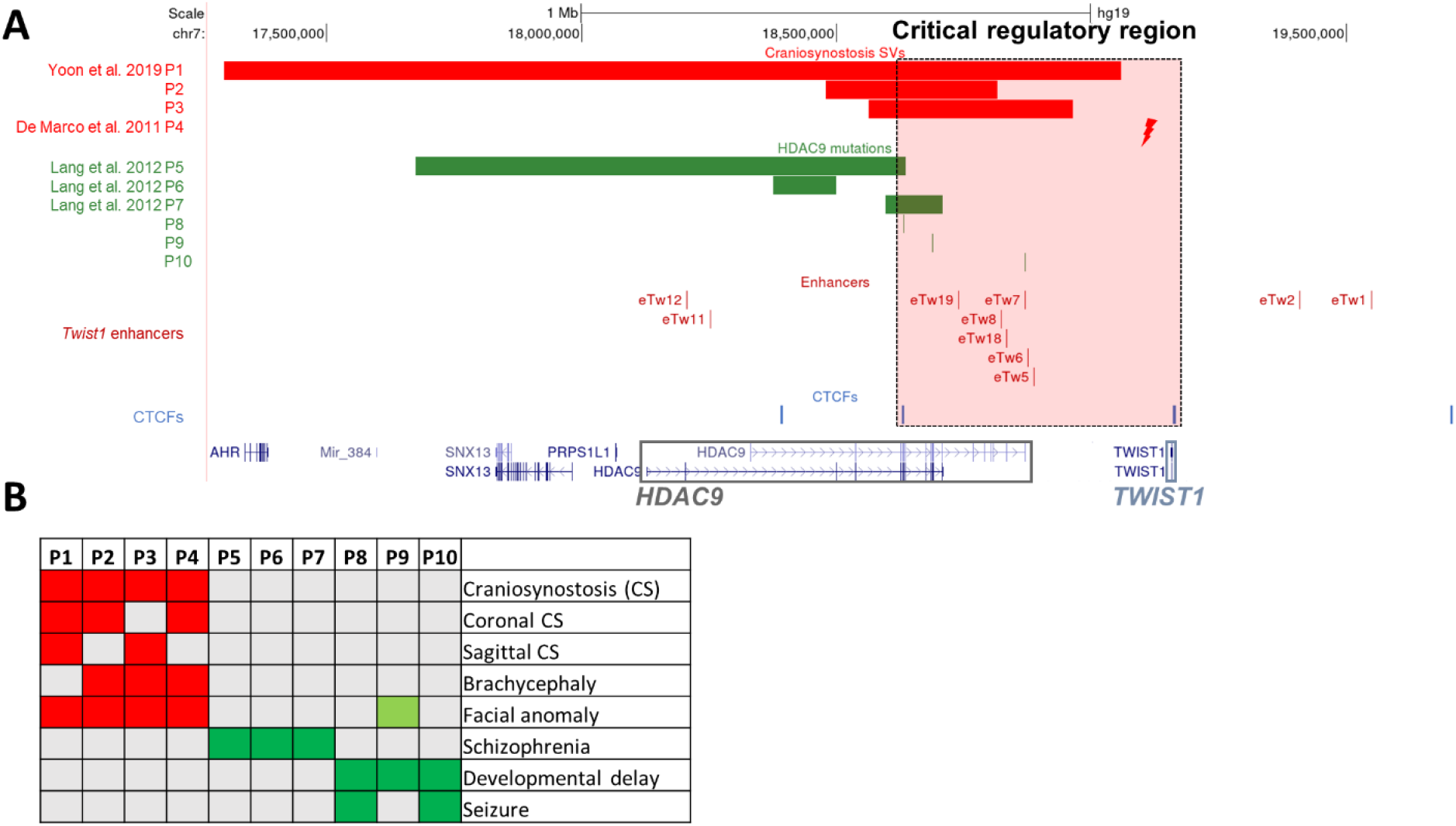
Structural variants containing *HDAC9* (but not *TWIST1*) in patients with craniosynostosis and neuronal disorders. **(A)** SVs encompassing *HDAC9* but not *TWIST1* indicate the potential location of a critical *TWIST1* regulatory region (highlighted by the dashed light pink rectangle). Red bars represent three craniosynostosis patients with *de novo HDAC9* deletions (P1-3) and red lightning represents translocation t(7;12)(p21.2;p12.3) breakpoint located between *HDAC9* and *TWIST1* in craniosynostosis patient. Green bars represent three schizophrenia patients (P5-7) with *de novo HDAC9* deletions and green lines represent three patients with neurological phenotypes (P8-10) with SNVs in *HDAC9*. Blue lines represent CTCF sites involved in chromatin looping in the *HDAC9-TWIST1* locus. Red lines represent *TWIST1* enhancers located in introns or exons of the *HDAC9* sequence and intergenic regions. **(B)** Human Phenotype Ontology heat map of patients’ common clinical features. Gray boxes represent either absent or unreported symptoms.

### Alteration of *Twist1* regulatory elements leads to a craniosynostosis-like phenotype

To evaluate the *in vivo* effect of aberrations within the *TWIST1* critical regulatory region on craniofacial and limb development, we investigated three mouse models. In the first two models, *Twist1* enhancers and Ctcf site residing in the *Hdac9* sequence were disrupted. Using the CRISPR/Cas9 genome editing system, we generated an **eTw5-7^Δ/Δ^** mouse model, homozygous for a deletion of three *Twist1* enhancers (eTw5-7) and four *Hdac9 exons* (exons 20-23, NM_001271386.1) **(Fig. 3A)**. In addition, we used a **Ctcf^Δ/Δ^** model^21^, in which the Ctcf binding site #5 involved in chromatin looping within the *Twist1* locus and located in the *Hdac9* sequence was deleted **(Fig. 3A)**. However, in both models (**eTw5-7^Δ/Δ^** and **Ctcf^Δ/Δ^**), the engineered deletions also encompass *Hdac9* protein-coding sequences, giving rise to the possibility that the Hdac9 protein is also involved in the phenotypic outcomes. To address this question, we generated a third mouse model carrying a large inversion (approximately 1Mb) of the entire *Hdac9* sequence that does not disrupt the *Hdac9* protein-coding sequence but repositions *Twist1* enhancers and potentially interferes with promoter-enhancer looping **(Fig. 3A)**.

**Figure 3.**
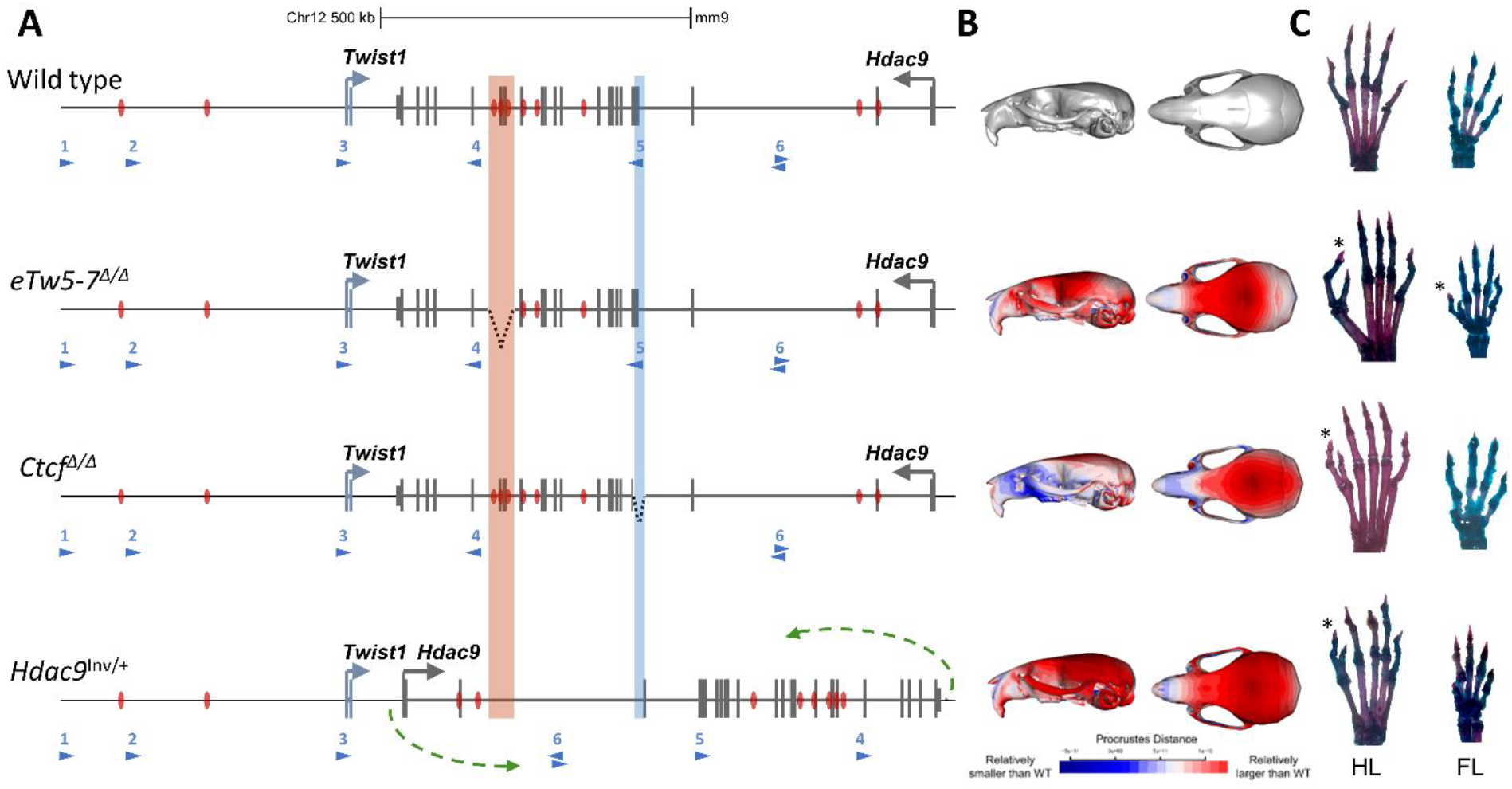
Alteration of *Twist1* regulatory elements in mice. **(A)** Scheme of the aberrations within the *Hdac9* sequence. In the eTw5-7^Δ/Δ^ model, a 23kb sequence, containing *Hdac9* exons 20-23 and *Twist1* enhancers (eTw5-7) was deleted (highlighted red rectangle). In the Ctcf^Δ/Δ^ model, a 1.5kb sequence containing *Hdac9* exons 6-7, was deleted (highlighted blue rectangle). In the *Hdac9*^INV/+^ model, the whole *Hdac9* sequence, 890kb long, was inverted. The inversion breakpoints are marked by green arrows (mm9, chr12:34,721,220-35,613,000). **(B)** Heat maps showing anatomical distributions of shape change compared to the wildtype for eTw5-7^Δ/Δ^, Ctcf^Δ/Δ^, and *Hdac9*^INV/+^ mice, demonstrated by side view (right) and superior view (left) (red is larger and blue is smaller compared to the grand mean). **(C)** Polydactyly was found in both hindlimb (71%) and forelimb (50%) of eTw5-7^Δ/Δ^ mice while polydactyly was found only in the hindlimb of Ctcf^Δ/Δ^ (32%) and *Hdac9*^INV/+^ (8%) mice.

Since TWIST1 haploinsufficiency leads to unilateral and bilateral coronal synostosis in both humans and *Twist1*^-/+^ mouse models, we tested whether deletions (eTw5-7^Δ/+^, eTw5-7^Δ/Δ^ and Ctcf^Δ/Δ^) or relocation (*Hdac9*^INV/+^) of *Twist1* regulatory elements have a significant impact on craniofacial development, including brachycephaly and craniosynostosis-like phenotype **(Fig. 3B)**. We used skeletal staining and micro-computed tomography (micro-CT) to quantify morphological effects on the skull. We analyzed adult (approximately 80 days of age) mouse skulls from our mouse models for *Twist1* regulatory elements (45 skulls of eTw5-7^Δ/Δ^, 18 skulls of eTw5-7^Δ/+^, 44 skulls of Ctcf^Δ/Δ^, 38 skulls of *Hdac9*^INV/+^) and compared them with a cohort of 25 wild-type littermates. We quantified 3D craniofacial shape from Micro-CT scan images using 68 standardized skeletal landmarks as in previous work **(Fig. S3)**^29^ using geometric morphometric methods. From the visualization of canonical variate analysis (CVA) and by comparing mean shapes among groups, we determined that phenotypic effects are not confined to a single feature but involve multiple regions of the skull. Our results indicate that deletion of *Twist1* enhancers (eTw5-7^Δ/Δ^) can cause a distinct set of alterations compared to wild-type morphology **(Movie S1-2)**. The first canonical variate (CV1) most clearly separates wildtype mice from eTw5-7^+/Δ^ and eTw5-7^Δ/Δ^ **(Fig. 4A)**, whereas the second canonical variate (CV2) demonstrates the morphological separation of wildtype from Ctcf^Δ/Δ^ and *Hdac9*^INV/+^ mice **(Fig. 4A, Movies S3-6)**. The one-way analysis of variance (ANOVA) model showed that skulls of the mouse models (eTw5-7^Δ/Δ^, Ctcf^Δ/Δ,^ and *Hdac9*^INV/+^) significantly differ from wild type skulls and their small skulls, recapitulating the skull phenotype of *Twist1*^-/+^ mice**^14^ (Figs. 3B and 4B)**. The Multivariate Analysis of Variance (MANOVA) showed significant shape differences among the different groups **(Table S4)**. Analyzing Procrustes (shape) distance for each individual from the wildtype mean shape, showed a significant difference in the average shape of eTw5-7^+/Δ^, eTw5-7^Δ/Δ,^ Ctcf^Δ/Δ,^ and *Hdac9*^INV/+^ mice from wildtype mice **(Fig. 4C)**. Using Alcian blue/Alizarin red staining, we found notable asymmetric in the cranial morphology of eTw5-7^Δ/Δ^ mice that resembles the unilateral and bilateral coronal synostosis found in humans with heterozygous *TWIST1* mutations **(Fig. 4D)**. To associate our mouse models to the uni- and bi-lateral craniosynostosis phenotype in humans, we quantified and compared asymmetry variances of the skulls using MANOVA for the asymmetric component of shape variation (the residuals of asymmetric Procrustes superimposition). The skulls of both eTw5-7^+/Δ^ and *Hdac9*^INV/+^ mice have significantly elevated asymmetry variances compared to wildtype **(Fig. 4E-F, Tables S5-6)**. However, no significant differences in cranial asymmetry were found between homozygous eTw5-7^Δ/Δ^ and Ctcf^Δ/Δ^ and the wildtype **(Fig. 4E-F, Tables S5-6)**. While all of our mouse models (e.g. both heterozygous and homozygous) showed significantly smaller skulls, only the heterozygous mice (e.g. eTw5-7^+/Δ^ and *Hdac9*^INV/+^) showed significant asymmetry of the skull, linking uni- and bi-lateral craniosynostosis in human to partial functional dose of Twist1 during craniofacial development. Overall, our results show that the uni- or bi-lateral craniosynostosis phenotype caused by Twist1 haploinsufficiency is due to disrupted *Twist1* regulatory elements and alteration of its spatiotemporal expression during development.

**Figure 4.**
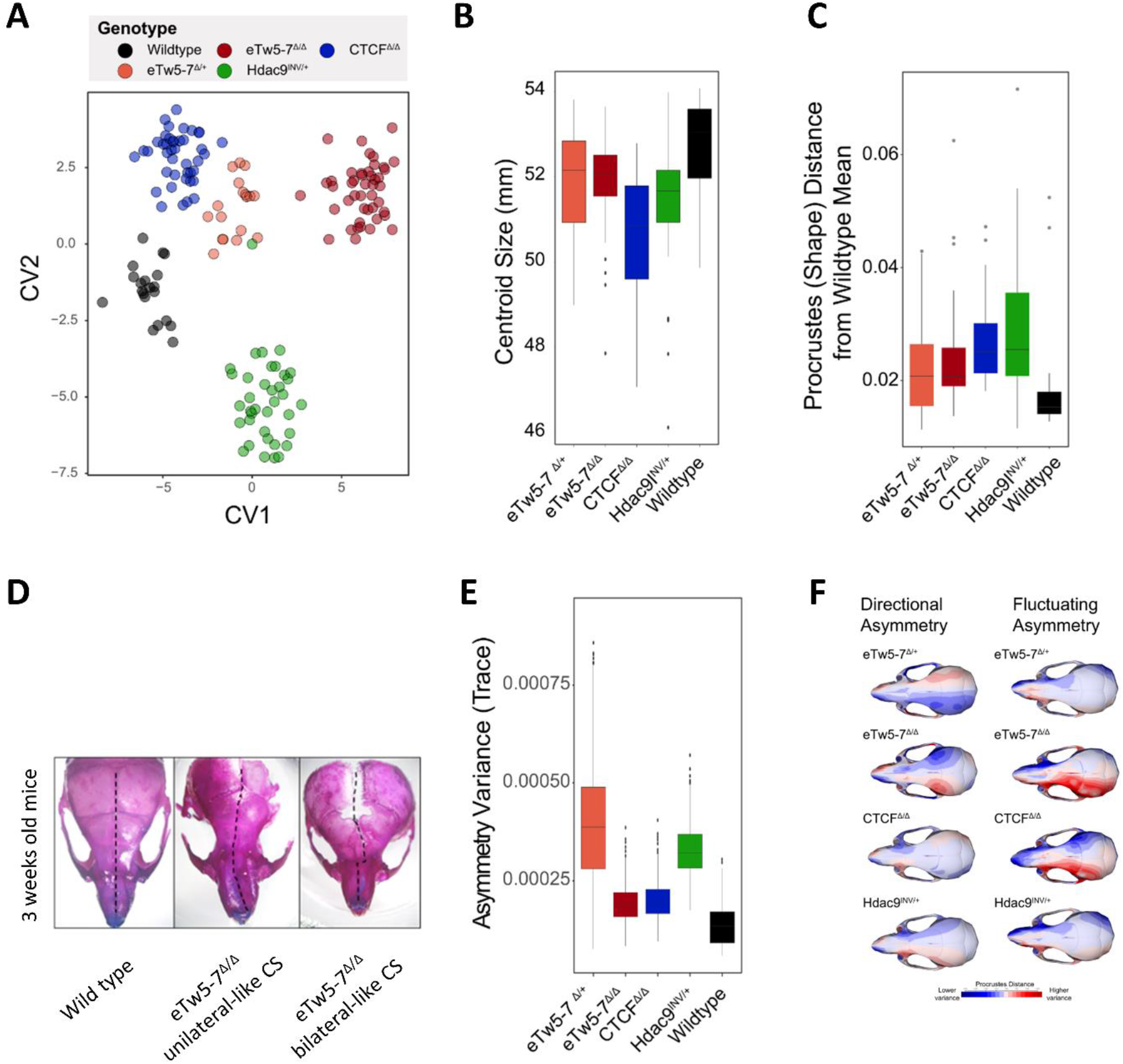
Deletions of *Twist1* regulatory elements lead to craniosynostosis-like morphology. **(A)** Canonical variate analysis (CVA) of micro-CT data from eTw5-7^Δ/Δ^, eTw5-7^Δ/+^, Ctcf^Δ/Δ^ and *Hdac9*^INV/+^ mice compared to wild-type. **(B)** Cranial size by genotype. **(C)** Boxplot for Procrustes (shape) distances for each individual from the wildtype mean shape. Note that since the Procrustes distance is unsigned, the average and minimum Procrustes distance for the wildtype is positive and not 0. **(D)** Representative pictures of stained skulls of 3 weeks old wildtype and eTw5-7^Δ/Δ^ mice, showing a range of variation in skull morphology. The dash black line represents the asymmetric structure with unilateral (middle) and bilateral (left) CS-like phenotypes. **(E)** Boxplot of the asymmetry variance differences among the different genotypes and the wildtype. It depicts group differences in the amount of variance explained by the asymmetric component of shape variation of the skull. **(F)** Heatmap visualizations for each asymmetry component (Directional Asymmetry and Fluctuating Asymmetry), displaying the patterns of shape variation in each genotype compared to the wildtype.

### Deletions of *Twist1* regulatory elements lead to pre-axial polydactyly

As Twist1 haploinsufficiency leads to limb malformations such as polydactyly, we evaluate the *in vivo* effect of disrupting *Twist1* regulatory elements on limb development using our mouse models. The eTw5-7^Δ/Δ^ mouse model showed high penetrance polydactyly in the hindlimb (71%) and forelimb (50%) **(Fig. 3C**). The polydactyly was observed in the right and/or left HL or FL, with 15% of the mice showing bilateral polydactyly. The Ctcf^Δ/Δ^ model^21^ showed hindlimb (HL) polydactyly with a similar penetrance (32%) as *Twist1*^-/+^ mice **(Fig. 3C)**. The polydactyly was observed in the right and/or left HL, with 20% of the mice showing bilateral polydactyly. Next, we evaluated whether *Twist1* is differentially expressed in our deleted *Hdac9* mouse embryos (eTw5-7^Δ/Δ^ and Ctcf^Δ/Δ^). Using qPCR and whole-mount *in-situ* hybridization, we discovered a reduction of *Twist1* expression in the anterior hindlimb bud of eTw5-7^Δ/Δ^ and Ctcf^Δ/Δ^ mice at E11.5 **(Fig. 5)**. These observations indicate that the dosage of Twist1 is critical and its haploinsufficiency can lead to partial penetrance and variable phenotypic expression. However, in these two models (eTw5-7^Δ/Δ^ and Ctcf^Δ/Δ^), the disrupted *Twist1* regulatory elements encompass the *Hdac9* protein-coding sequences, suggesting that the lack of Hdac9 protein could also contribute to the phenotype. We showed that *Hdac9* is not expressed in either wildtype or eTw5-7^Δ/Δ^\Ctcf^Δ/Δ^ mice at the E11.5 limb bud indicating that Hdac9 does not contribute to the polydactyly phenotype **(Fig. 5C)^8^**. Moreover, we showed that while homozygosity for the inversion (*Hdac9*^INV/INV^) is lethal as in *Twist1* null mice, the heterozygous mice (*Hdac9*^INV/+^) are viable and showed HL polydactyly (8%, **Fig. 3C**). We found that *Twist1* levels were reduced in HL of *Hdac9*^INV/+^ embryos compared to wild types **(Fig. S4).** Furthermore, no reduction in Hdac9 protein level was found in the cortical brain (where *Hdac9* is highly expressed) compared to adult control mice **(Fig. S5)**. Our results imply that the polydactyly phenotype of our mouse models is due to the alteration of *Twist1* regulation and not the Hdac9 protein.

**Figure 5:**
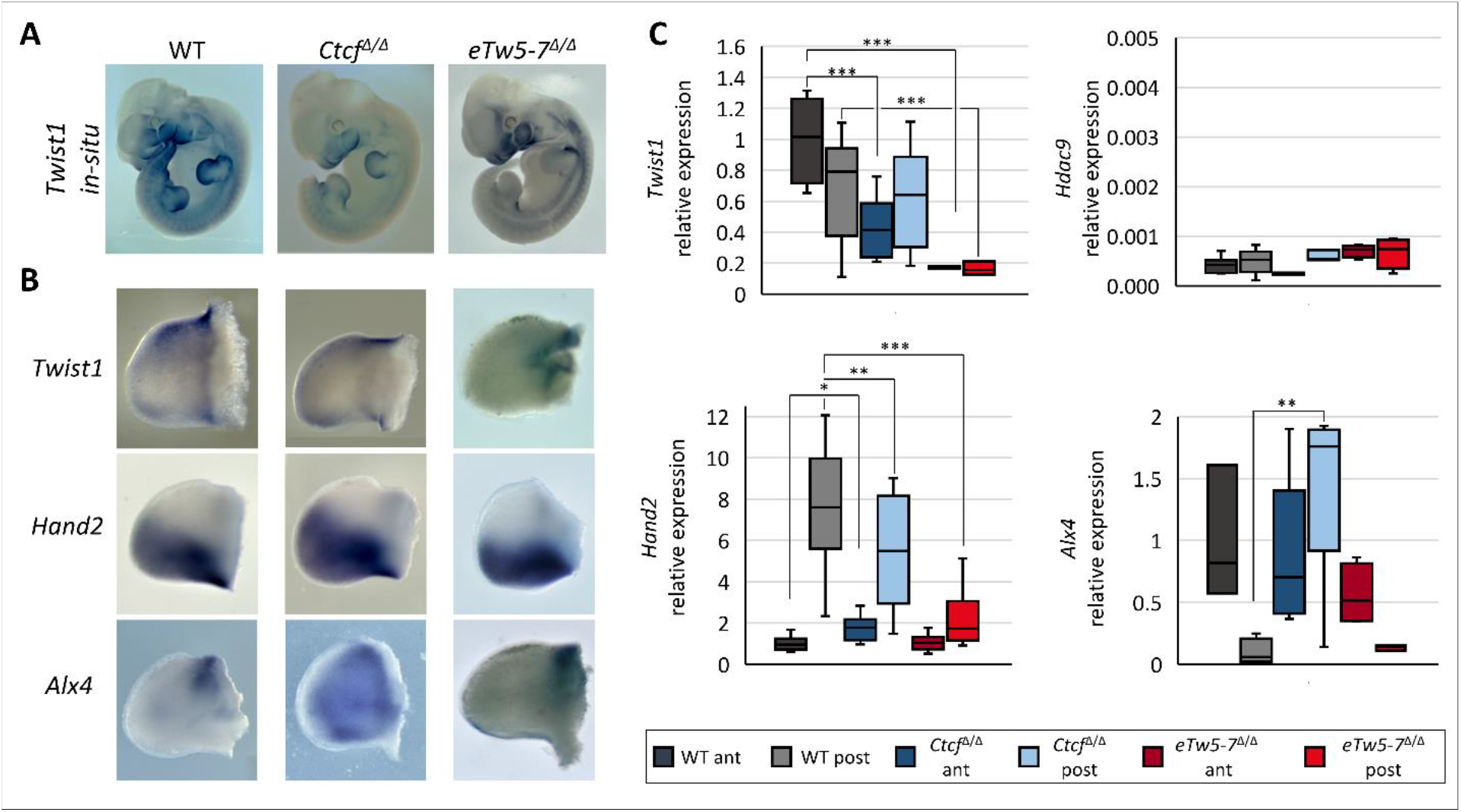
Deletions in *Hdac9* that alter *Twist1* regulatory elements lead to dis-regulation of Shh pathway genes. **(A)** Whole-mount *in-situ* hybridization of E11.5 mouse embryos showing reduced *Twist1* expression pattern in eTw5-7^Δ/Δ^ and Ctcf^Δ/Δ^. **(B)** Whole-mount *in-situ* of E11.5 hindlimb mouse embryos for *Twist1* and its target *Shh* pathway genes, *Hand2* and *Alx4. Twist1* expression is reduced especially in the anterior limb bud. *Hand2* and *Alx4* expression along the anterior/posterior axes is lost in the mutants. **(C)** Quantitative real-time PCR analyses in E11.5 HL buds show a significantly decreased *Twist1* expression in anterior limb buds of Ctcf^Δ/Δ^ (*P*= 9×10^-4^) and in the entire limb bud of eTw5-7^Δ/Δ^ (*P*=5×10^-4^). No expression of *Hdac9* in the limb buds of wildtype, Ctcf^Δ/Δ^ or eTw5-7^Δ/Δ^ embryos. Significant decrease of *Hand2* expression in posterior limb buds of Ctcf^Δ/Δ^ (*P*=3×10^-2^) and eTw5-7^Δ/Δ^ (*P*=1×10^-5^) embryos. Significant increase of *Alx4* expression in posterior limb buds of Ctcf^Δ/Δ^ embryos (*P*=8.8×10^-3^).

### Deletion of *Twist1* regulatory elements alters the expression of Shh pathway genes

To evaluate the effect of altered regulatory elements on *Twist1* target genes, we analyzed the expression of *Hand2, Ptch1, Alx4* and *Gli3*, known Twist1 target genes in the Shh pathway, both directly and indirectly. These transcription factors have unique and specific expression patterns along the anterior-posterior (A-P) axis of the limb bud, required for digit number and identity. Indeed, we showed that disrupted *Twist1* regulatory elements not only reduced the expression of *Twist1* but affect the spatiotemporal expression of its target genes along the A-P axis **(Fig. 5B-C)**. Using qPCR, we observed that the expression pattern of *Hand2* was reduced in the posterior limb bud of eTw5-7^Δ/Δ^ and Ctcf^Δ/Δ^ (P<0.05; **Fig. 5**). Moreover, using *in-situ* hybridization, *Hand2* expression extended beyond the posterior boundary and diffused towards the anterior domain. This ectopic Hand2 expression led to a disproportion of the A-P pattern in the limb bud **(Fig. 5B-C)**. *Ptch1*, a gene coding for transmembrane Shh receptor that is differentially expressed in A-P axes, also lost its A-P limb bud expression in eTw5-7^Δ/Δ^ **(Fig. S6)**. Furthermore, consistent with the negative feedback loop of Shh regulators, Shh antagonists *Alx4* and *Gli3* are both lacking precise anterior expression patterns and are extended towards posterior domains in Ctcf^Δ/Δ^ and eTw5-7^Δ/Δ^ embryos **(Fig. 5B-C, Fig. S6)**. *Alx4* expression levels are significantly increased in the posterior limb buds *and Gli3* expression level is reduced in the anterior limb buds of Ctcf^Δ/Δ^ embryos **(Fig. 5B-C, Fig. S6)**. Therefore, deletions of the eTw5-7 enhancers or Ctcf site led to a significant reduction of *Twist1* expression levels, and ectopic expression of *Hand2 and Alx4*. These Shh pathway genes are no longer differentially expressed along the A-P axis and have a diffused pattern that likely led to inconsistency of Shh pathway signals and eventually resulted in high penetrance polydactyly.

### Deletion of *Twist1* enhancers and a Ctcf site affects chromatin looping at the *Twist1-Hdac9* locus

To elucidate the effect of *Twist1* regulatory element deletions on chromatin organization within the *Twist1-Hdac9* locus, we performed UMI-4C on E11.5 forelimbs, hindlimbs, and branchial arches. We compared the interaction maps of wildtype, eTw5-7^Δ/Δ^, and Ctcf^Δ/Δ^ embryos using the *Twist1* promoter as a viewpoint **(Fig. 6)**. The *Twist1* promoter displayed frequent interactions with a region that contains the cluster of three enhancers, *eTw-5, 6 and 7* in limb buds and the branchial arch, with the highest interaction frequency being observed in the limb buds **(Fig. 6)**. These three enhancers were characterized with limb enhancer activity and two of them (*eTw-5, 6*) were also active in the branchial arches^8^. In addition, the *Twist1* promoter region was also involved in frequent interactions with two upstream Ctcf sites (Ctcf#1 and Ctcf#2) and two downstream Ctcf sites (Ctcf#5 and Ctcf#6), both in limb buds and branchial arch **(Fig. 6B)**. While the *Twist1-Hdac9* locus contains several Ctcf sites, these four Ctcf-bound interacting regions are occupied by Ctcf in the limb bud at E1 1.5^24^, supporting their role in the 3D chromatin organization of the locus, facilitating Twist1 regulatory interactions **(Fig. 6B)**. As expected, in both eTw5-7Δ/Δ and CtcfΔ/Δ embryos, interactions between the Twist1 promoter and the targeted region (eTw5-7 and Ctcf, respectively) were severely reduced **(Fig 6B-C, Fig. S7)**. In eTw5-7^Δ/Δ^ embryos, the *Twist1* promoter region showed a 5.9-fold reduction in interaction frequency with the eTw5-7 region in HL and a 3.4-fold reduction in FL (P=1.55×10^-62^, 1.42×10^-22^, respectively). In Ctcf^Δ/Δ^ embryos, there was a 3.2-fold reduction in interaction frequency with the Ctcf region in HL (P=1.42×10^-11^) and a 2.8-fold reduction in FL (P=1.97×10^-8^). Intriguingly, we also observed reduced interactions of the Twist1 promoter with the Ctcf site in eTw5-7^Δ/Δ^ embryos and vice versa in Ctcf^Δ/Δ^. In eTw5-7^Δ/Δ^ embryos, the deletion of the eTw5-7 enhancer cluster led to a 1.7-fold reduction in interaction frequency with the Ctcf region in HL (P=0.05) and a 1.2-fold reduction in the FL (P>0.05) **(Fig. 6C)**. In Ctcf^Δ/Δ^ embryos, the deletion of the Ctcf site altered the interaction frequency with the eTw5-7 region, with a 1.4-fold reduction of interaction frequency detected in the HL (P=5.42×10^-8^) and a 1.3-fold reduction in FL (P= 6.39×10^-5^) (**Fig. 6D)**. These interaction profiles imply that both the eTw5-7 enhancer cluster and the Ctcf site are partially dependent on each other for interaction with the Twist1 promoter. Similar interaction profiles were found in E11.5 BA **(Fig. 6B)**. In eTw5-7^Δ/Δ^ embryos, the *Twist1* promoter region showed a 2.6-fold loss of interactions with the eTw5-7 region (P=1.3×10^-19^), but no significant change was observed at the Ctcf site in the BA **(Fig. 6C)**. Furthermore, in Ctcf^Δ/Δ^ embryos, the *Twist1* promoter region showed a 3-fold loss of interactions with the Ctcf region (P=1.4×10^-6^), but no significant change was observed at the eTw5-7 region **(Fig. 6D)**. This suggests that while the chromatin looping at *the Hdac9-Twist1* locus is similar in both limb bud and BA, the interaction frequencies and the dependency of the regulatory elements involved in *the Twist1* transcriptional mechanism are different. Overall, the *Twist1* promoter region along with enhancers and Ctcf binding sites compose a regulatory unit, likely functioning together to execute precise *Twist1* expression during limb and craniofacial development.

**Figure 6:**
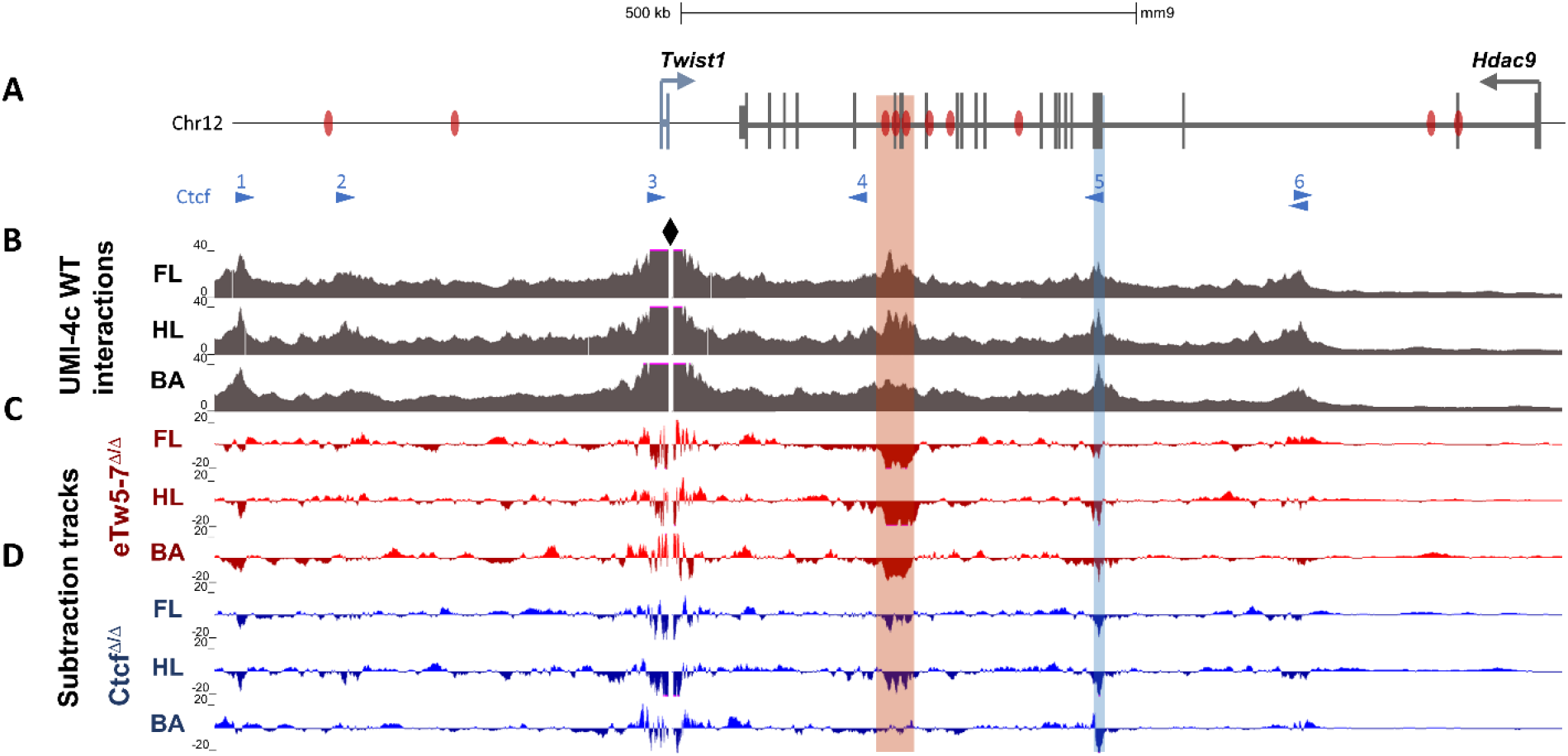
Deletions of *Twist1* regulatory regions affect chromatin looping in the *Hdac9-Twist1* locus. **(A)** A scheme of the Hdac9-Twist1 locus with Twist1 enhancers marked by black bars and Ctcf-bound sites^24^ (and motif directionality) marked by blue triangles. **(B)** UMI-4C tracks (merged from two biological replicates) of WT E11.5 forelimb (FL), Hindlimb (HL), and Branchial arch (BA) demonstrate that *Twist1* interaction patterns are largely conserved between limb buds and branchial arch. The *Twist1* promoter serves as a viewpoint and is highlighted by the black diamond shape. Targeted eTw5-7 and Ctcf regions are indicated by red and blue rectangles, respectively. **(C-D)** Subtraction tracks showing differential *Twist1* promoter interactions in FL, HL, and BA of eTw5-7^Δ/Δ^ and Ctcf^Δ/Δ^ embryos compared to WT (see also Fig. S7).

## Discussion

In this study, we deciphered the role of SVs at the *HDAC9-TWIST1* locus in craniosynostosis and craniofacial malformations. We identified several chromosomal aberrations affecting *HDAC9* but not *TWIST1* in patients with craniosynostosis. This suggests that the disruption of a critical *TWIST1* regulatory region, within the *HDAC9* sequence, leads to craniofacial malformations. We initially characterized 6 transcriptional enhancers located in the *HDAC9-TWIST1* synteny block that regulates spatiotemporal *Twist1* activity during craniofacial development. These enhancers also reside in regions that interact with the Twist1 promoter in BA and limb bud of E11.5 mouse embryos. Each enhancer has a discrete activity pattern that recapitulated aspects of *Twist1* expression during development. The partially overlapping activity pattern of the analyzed *Twist1* enhancers might ensure the robustness of *Twist1* expression during craniofacial and limb development. For example, *eTw-5, eTw-18*, and *eTw-19* are active enhancers in zebrafish BA1-7, while *eTw-2* and *eTw-6* enhancers are active in BA1-2, and eTw11 enhancer is active in BA3-7, indicating that the activity of each enhancer, along with the overlapping activity between enhancers, is essential for proper spatiotemporal *Twist1* expression.

Loss of function of craniofacial and limb *Twist1* enhancers and Ctcf sites emphasizes the crucial role of *TWIST1* transcriptional regulation on the penetrance and expression level of the craniosynostosis and limb phenotype. All three models (eTw5-7^Δ/Δ^, Ctcf^Δ/Δ^, and *Hdac9*^INV/+^) showed a skull phenotype but the effect on the skull shape depended on the characteristics of the disrupted regulatory elements **(Fig. 4E-F)**. For example, both eTw5-7^Δ/Δ^ and Ctcf^Δ/Δ^ showed small centroid size but with a different effect on skull shape **(Fig. 4, Movie S1)**. In addition, we found significant asymmetry in the skulls of both eTw5-7^+/Δ^ and *Hdac9*^INV/+^ mice that resembles uni-lateral craniosynostosis, while homozygous mice show small skull with no significant asymmetry, that resembles bi-lateral craniosynostosis **(Fig. 4E-F)**. The partial penetrance and variable expression of the phenotype are also emphasized in the developing limb. While homozygous eTw5-7^Δ/Δ^ mice showed both hindlimb and forelimb polydactyly with high penetrance (77%, **Fig. 3**) and variable expression (right and/or left hind\fore limb), heterozygous eTw5-7^Δ/+^ mice have no limb phenotype. Similar observations were made for homozygous Ctcf^Δ/Δ^ mice, which showed hindlimb polydactyly with variable expression (right and/or left hindlimb) and partial penetrance (32%) as seen in *Twist1*^-/+^ mice **(Fig. 3)**. The phenotypic similarity of *Twist1* null mouse was also observed for the homozygous *Hdac9* inversion mice, which are lethal, while the heterozygous mice (Hdac9^INV/+^) are viable and show hindlimb polydactyly (10%, **Fig. 3**). In addition, the deletions of eTw5-7 enhancers or the Ctcf site led to a significant reduction of *Twist1*, specifically in anterior limb buds, and ectopic expression of *Twist1* target genes, including *Hand2, Alx4* as well as *Ptch1* and *Gli3*, respectively **(Fig 5, Fig. S6)**. These Shh pathway genes are no longer differentially expressed along the A-P axis and have a diffused expression pattern without significant difference between anterior and posterior of the limb bud, which likely led to inconsistency of Shh pathway signals and eventually resulted in variable penetrance polydactyly. Indeed, each of our mouse models shows a specific disruption of *Twist1* expression, emphasizing the role of *Twist1* regulatory elements in finetuning spatiotemporal *Twist1* expression. Overall, the type and number of disrupted regulatory elements can be associated with uni- and bi-lateral (or none) craniosynostosis-like phenotype, as well as polydactyly based on a threshold dose of Twist1.

The tissue-specific activities of these craniofacial and limb enhancers are supported by specific chromatin interactions within the *Twist1-Hdac9* locus in the limb bud and branchial arch. UMI-4C data demonstrated that the *Twist1* promoter region interacts with *eTw-5, 6, 7* and 18 enhancers in the limb bud and branchial arch. Interestingly, the *Twist1* promoter region that contains a Ctcf site, interacts with four distal Ctcf-bound sites, which together enabling 3D chromatin interactions required for *Twist1* transcriptional regulation **(Fig. 6)**. Indeed, reduced chromatin looping was observed upon deletion of the *eTw-5-7* and Ctcf sites **(Fig. 6)**. Interestingly, in limb buds, *Twist1* interaction frequency with both sites appeared to be co-dependent. This suggests that the Twist1 promoter region along with enhancer elements and architectural Ctcf-bound sites compose a regulatory unit, functioning in unison to finetune precise *Twist1* expression.

In summary, SVs represent an important mutational force that shapes genome function and contributes to germline and somatic diseases. As SVs are responsible for ~25% of all rare protein-truncating events per genome, they have a major effect on protein sequences but also on cis-regulatory elements that control spatiotemporal gene expression. Here, we showed that SVs that affect the HDAC9 sequence can also modulate basic mechanisms of gene regulation controlling the expression of a nearby gene, *Twist1*. These SVs affect *Twist1* regulatory elements and disrupt higher-order chromatin organization, leading to a phenotype that is not associated with the HDAC9 protein. Thus, careful interpretation is required when considering the molecular basis of SVs identified in human patients.

## Materials and Methods

### Ethics Statement

DNA samples were obtained from all available samples following informed consent and approval of the Soroka Medical Center Internal Review Board (IRB). Clinical phenotyping was determined by an experienced pediatric and geneticist.

All animal work was approved by the Ben Gurion Institutional Animal Care and Use Committee protocol number 52-09-2016.

### Generating Hdac9\Twist1 mouse models

Two mouse strains, eTw5-7^Δ/Δ^ and *Hdac9* ^Inv/+^, were created using a modified CRISPR/Cas9 protocol^30^. In addition, we exploited a mouse model where exons 6 and 7 of the Hdac9 sequence, along with a Ctcf site at the interionic sequence were deleted^21^. Briefly, for eTw5-7^Δ/Δ^, two sgRNAs targeting a 23 kb sequence that encompasses the three enhancers and exons 17–20 of *Hdac9* (23,137bp; chr12:34883772–34906909; mm9) were designed using CHOPCHOP^31^. Similarly, for *Hdac9* inversion two gRNAs were designed, targeting regions delimitating the whole Hdac9 gene (chr12:34721220-35613000, 891,781 bp). No potential off-targets were found when searching for matches in the mouse genome (mm9) when allowing for up to two mismatches in the 20 nucleotide-long sequence preceding the PAM sequence. The T7 promoter was added to the sgRNA template, and the whole cassette was chemically synthesized by IDT. The PCR amplified T7-sgRNA product was used as a template for *in vitro* transcription using the MEGAshortscript T7 kit (Thermo Fisher Scientific, USA). Cas9 mRNA was transcribed *in vitro* using the mMESSAGE mMACHINE T7 kit (Thermo Fisher Scientific, USA). The DNA template for *in vitro* transcription containing the humanized *Streptococcus pyogenes* Cas9 gene was PCR-amplified from the px330 plasmid. eTw5-7^Δ/Δ^ and *Hdac9*^INV/+^ mice were generated by injecting a mix of Cas9 mRNA (final concentration of 100 ng/ul) and sgRNA (50 ng/ul) in injection buffer (10 mM Tris, pH 7.5; 0.1 mM EDTA) into the cytoplasm of C57black embryos in accordance with the standard procedure approved by Ben-Gurion University. Female mice of the ICR (CD-1) strain were used as foster mothers. F0 mice were genotyped using PCR to detect the deletion of enhancers from the mouse genome **(Table S7)**.

### Genomic data analyses

Public available ChIP-seq and ATAC-seq data sets of E10.5/E11.5 mice embryos craniofacial tissues (i.e. Mx, Md, PA2, FNP) which used enhancer-associated marks (EP300, H3K27ac)^23,32^ as well as human embryonic ChIP-seq data for chromatin modifications (H3K4me1, H3K4me2, H3K4me3, H3K36me3, H3K27ac, H3K27me3) from stages CS13-CS15 on craniofacial tissues ^22^ were obtained and analyzed for redundancy, as well as evolutionary conservation. Overlapping peaks were defined as when at least 1 bp regions overlapped.

### Transgenic zebrafish enhancer assay

Primers were designed to amplify the candidate enhancer sequences from human genomic DNA (Table S7). PCR products were cloned into the E1b-GFP-*Tol2* enhancer assay vector containing an E1b minimal promoter followed by the gene for GFP^33^ These constructs were injected into zebrafish embryos using standard procedures. For statistical significance, at least 100 embryos were injected per construct in at least two different injection experiments along with *Tol2* mRNA to facilitate genomic integration^34^. GFP expression was observed and annotated 48 and 72 hpf. An enhancer was considered as a positive enhancer when at least 30% of the live embryos showed a consistent GFP expression pattern.

### Craniofacial Morphometric Analysis and Statistical Analysis

Mice were scanned using a Scanco VivaCT scanner at 35 micron resolution. The total sample size for this analysis was 170 skulls (25 Wildtype, 38 Hdac^Δ/+^, 44 CTCF^Δ/Δ^, 18 eTw5-7^Δ/+^ and 45 eTw5-7^Δ/Δ^). To quantify craniofacial shape, we used the standard set of 68 3D landmarks used in previous work ^35^,^36^. We used geometric morphometric methods to perform quantitative analyses to statistically evaluate and visualize patterns of variation in craniofacial shape and size, using the software R. We performed a Generalized Procrustes Superimposition Analysis (GPA) to extract the aligned Procrustes shape coordinates from the 3D landmark data, using the R package *geomorph*^37^. We quantified cranial size as the centroid of each landmark configuration, which is calculated as the square root of the sum of the squared distances from each landmark to the landmark set centroid for each specimen, using *geomorp*. We fitted a Multivariate Analysis of Variance (MANOVA), implemented in the geomorph R package^37^, to determine differences in craniofacial size and shape between the different genotypes and the wildtype group.

To quantify shape differences between each genotype and the wildtype, we calculated the shape (Procrustes) distance for each individual in the sample to the wildtype mean. This generates a distribution of distances for each genotype including the wildtype. Since no individual is identical to its group genotype mean and because the Procrustes distance is unsigned, the mean shape distance of wildtype individuals from the wildtype mean is positive and not zero. To visualize the shape effects that distinguish each genotype from the wildtype mean, we determined the vector of Procrustes coordinate differences between each genotype mean and the wildtype mean. This vector was then used to generate a heatmap showing the anatomical distribution of these distances and to generate 3D morphs. Since the mean differences themselves are fairly subtle and difficult to see on 3D morphs, we can exaggerate them by multiplying the shape difference vector by an arbitrary constant. This method was used to generate exaggerated versions of the genotype effects in Figure 3 and the 3D morph movies in the supplemental data.

To assess craniofacial asymmetry, we decomposed skull shape variation and extracted directional asymmetry (DA) and fluctuating asymmetry (FA) data. Directional asymmetry (DA) is the pattern of consistent shape variation between left and right sides, involving developmental mechanisms and/or differential gene expression^38^, and the DA component for each individual is calculated as the shape difference between sides (left and right). Fluctuating asymmetry (FA) captures small asymmetric shape differences between left and right sides that, unlike DA, does not follow a pattern and is displayed as random or residual shape variation ^38^. A higher amount of shape variance explained by the FA component within the asymmetric shape component can be associated with developmental instability ^39 40^. The FA component is calculated for each specimen and each side (left and right) as the adjusted deviation for the mean DA.

To visualize shape changes among groups and variation along the axes that distinguish groups, we performed canonical variate analysis (CVA) as well as principal components analysis (PCA), using the R packages *geomorph* ^37^ and *morpho* ^41^. The vector displacements from these analyses were used to visualize shape variation using morphs of 3D meshes or as heatmaps, using a thin-plate spline method to calculate distances between the reference mesh and the target mesh. Negative values (inside the reference mesh) were visualized with red colors, whereas positive values (outside the reference mesh) were visualized with blue color values. Distances close to 0 (target mesh practically in the same position as the reference mesh) were visualized as white. Both the morphs and heatmaps displaying differences between wildtype (reference) and each genotype (target) were generated using the wildtype average mesh and wildtype landmark coordinates and the average shape coordinates for each genotype (**Fig. 3B**). Morphs and heatmaps from the CVA analyses were generated from the average mesh and average landmark coordinates and the shape coordinates of the extreme ends of the CVA axes (CV1-2). Importantly, the determination of the statistical significance of differences among groups is based on the MANOVA and not on the CVA analyses, which were only performed to visualize the patterns of shape variation found in the MANOVA^42^.

### Gene expression analyses

Mouse E10.5 and E11.5 limb buds (whole or dissected for anterior/posterior), as well as adult brains, were dissected. Total RNA was isolated using a Total RNA purification micro kit (NORGEN, Cat. 35300, Canada) according to the manufacturer’s protocol. Next, cDNA was synthesized using 5X All-In-One master mix (abm, Cat. G592, USA). qPCR was performed using KAPA SYBR^®^ FAST qPCR Master Mix (Sigma Aldrich) and run on the C1000 Touch™ thermal cycler (BIO-RAD, USA). Samples were tested in duplicates. The specificity and absence of primer dimers were controlled by denaturation curves. β-Actin (*Actb*) mRNA was used for normalization.

### Whole-mount *In-Situ* hybridization

Mouse E10.5 and E11.5 embryos were fixed in 4% paraformaldehyde. Vectors containing mouse *Twist1, Hand2* and *Alx4* were used as templates for digoxygenin-labeled probes. Mouse whole-mount *in situ* hybridizations were performed according to standard procedures^43^.

### UMI 4C

UMI-4C was performed according to the protocol by Schwartzman et al.^44^. In brief, limb bud and branchial arch tissues of mouse E11.5 embryos were micro-dissected, dissociated into single cells, and crosslinked for 10 min using 2% formaldehyde. Crosslinked cell pellets were snap-frozen and stored at −80°C until further processing^44^. Frozen pellets were resuspended in 250μL pre-diluted DpnII buffer, 7.5μL pre-heated 10% SDS and incubated on a thermomixer for 1h at 37°C, shaking at 900 RPM. After adding 75μL 10% Triton X-100 the solution was incubated again (1h, 37°C, 900 RPM). The chromatin was digested using 300U DpnII (NEB, R0543L) in three stages (100U for 2h, 100U overnight, 100U for 2h) at 37°C and 900 RPM. The solution was incubated at 65°C for 20 min to inactivate the restriction enzyme and put on ice. Next, the chromatin was ligated by adding 2000U of T4 DNA ligase (NEB, M0202M) and 10x T4 DNA ligase buffer to a total volume of 650μL and incubating the solution overnight at 16°C and 300 RPM. Ligated chromatin was de-crosslinked by incubation with 4μL proteinase K (20mg/mL, Qiagen, Cat. 19131, Germany), overnight for 65°C and 300 RPM). A 3C template was then purified using 1x Ampure XP beads (Beckman Coulter, A63881, USA). Up to 4μg 3C template per sample was sheared using microTUBE snap-cap tubes (Covaris, 520045) in a Covaris M220 sonicator to an average fragment length of 300bp. UMI-4C sequencing libraries were generated using the NEBNext Ultra II library prep kit (NEB, E7645L). For each sample, library prep was performed in four parallel reactions with a maximum input of 1000ng per reaction, including a size selection targeting fragments with a length of 300-400bp (unless the input was <100ng) and 4-8 cycles of PCR enrichment depending on the input. Next, two nested PCR reactions were performed to enrich for fragments captured by the viewpoint of interest, both using 2μL 10mM Illumina enrichment primer 2 (CAAGCAGAAGACGGCATACGA) and either a viewpoint-specific “upstream” (reaction 1, 2μL 10mM, CTGTGACAGCAGTAGTGGCA) or “downstream” (reaction 2, 2μL 10mM, AATGATACGGCGACCACCGAGATCTACACTCTTTCCCTACACGACGCTCTTCCGATCTCTTCGACGCTCTGGGTGAT) primer. For each sample, we performed up to 8 nested PCR reactions in parallel with an input of 100-200ng per reaction using the KAPA2G Robust ready mix (Sigma-Aldrich, KK5702) in a volume of 50μL. PCR program: 3 min 95°C, 20 cycles (18 cycles for reaction 2) of 15 sec 95°C, 15 sec 55°C, 60 sec 72°C and final elongation of 5 min 72°C. Between PCR reactions the product was cleaned up using 1x AmpureXP beads and eluted in 21μL. The final PCR product was cleaned up using 0.7x AmpureXP beads and eluted in 25μL. Reactions per sample were pooled and library concentration was quantified via qPCR using the KAPA SYBR FAST qPCR Master Mix (Sigma-Aldrich, KK4602). UMI-4C libraries were pooled and sequenced on an Illumina HiSeq or NovaSeq (paired-end, 2×150 cycles). Reads were first filtered based on the presence of the downstream primer sequence (20% mismatch allowed). The resulting fastq files were then used as input to the UMI-4C R package (https://github.com/tanaylab/umi4cpackage) to generate genomic interaction tracks, representing UMI counts (i.e. unique interactions) per genomic restriction fragment. The package was then used to generate smoothed, viewpoint-specific interaction profiles for the region of interest. For each profile, interaction counts were normalized to the total UMI count within the profile. Normalized profiles were subtracted to identify differentially interacting regions. Statistical significance of differential interactions was tested for two loci (“eTw5-7”: chr12:34883878-34906858, “Ctcf”: chr12:35116684-35117991) using a Chi-square test within the UMI-4C R package (p4cIntervalsMean function).

### Alcian blue/Alizarin red staining

Skulls of three weeks old mice were processed and stained for bone and cartilage as previously described^45^. Mice were euthanized and their skulls were dissected. The skin, eyes, organs, and adipose tissue were removed using forceps. Skulls were then fixed by two overnight changes of 95% ethanol, followed by two days in acetone. Next, the cartilage was stained using alcian blue staining solution (0.03 % (w/v), 80 % EtOH, 20 % (glacial) acetic acid) for two days. Then, skulls were distained and postfixed by two overnight changes of 95% ethanol and then cleared in 1% KOH overnight in 4°_C_, followed by bone staining with alizarin red (0.005 % (w/v) in 1 % (w/v) KOH) for three days. Finally, specimens were cleared using 1% KOH and stored in 100% glycerol.

## Acknowledgments

This research was supported by the Israel Science Foundation (https://www.isf.org.il). NH and RYB were partially supported by this grant. This work was partly supported by the Research Foundation Flanders (FWO) under grant G044615N and 1520518N. In addition, E.D. and S.V. were respectively supported by a doctoral and postdoctoral fellowship of the FWO Research Fund.

## Notes

### Competing Interest Statement

The authors have declared no competing interest.

## References

1. Collins, R.L. et al. A structural variation reference for medical and population genetics. Nature 581, 444–451 (2020).

2. Audano, P.A. et al. Characterizing the Major Structural Variant Alleles of the Human Genome. Cell 176, 663–675 e19 (2019).

3. Spielmann, M., Lupianez, D.G. & Mundlos, S. Structural variation in the 3D genome. Nat Rev Genet 19, 453–467 (2018).

4. Lupianez, D.G. et al. Disruptions of topological chromatin domains cause pathogenic rewiring of gene-enhancer interactions. Cell 161, 1012–25 (2015).

5. Birnbaum, R.Y. et al. Coding exons function as tissue-specific enhancers of nearby genes. Genome Res 22, 1059–68 (2012).

6. Birnbaum, R.Y. et al. Systematic dissection of coding exons at single nucleotide resolution supports an additional role in cell-specific transcriptional regulation. PLoS Genet 10, e1004592 (2014).

7. Hirsch, N. & Birnbaum, R.Y. Dual Function of DNA Sequences: Protein-Coding Sequences Function as Transcriptional Enhancers. Perspect Biol Med 58, 182–95 (2015).

8. Hirsch, N. et al. Unraveling the transcriptional regulation of TWIST1 in limb development. PLoS Genet 14, e1007738 (2018).

9. Birnbaum, R.Y. et al. Functional characterization of tissue-specific enhancers in the DLX5/6 locus. Hum Mol Genet 21, 4930–8 (2012).

10. Qin, Q., Xu, Y., He, T., Qin, C. & Xu, J. Normal and disease-related biological functions of Twist1 and underlying molecular mechanisms. Cell Res 22, 90–106 (2012).

11. Zhang, Z. et al. Preaxial polydactyly: interactions among ETV, TWIST1 and HAND2 control anterior-posterior patterning of the limb. Development 137, 3417–26 (2010).

12. Cho, E. et al. Saethre-Chotzen syndrome with an atypical phenotype: identification of TWIST microdeletion by array CGH. Childs Nerv Syst 29, 2101–4 (2013).

13. Miller, K.A. et al. Diagnostic value of exome and whole genome sequencing in craniosynostosis. J Med Genet 54, 260–268 (2017).

14. Parsons, T.E. et al. Craniofacial shape variation in Twist1+/- mutant mice. Anat Rec (Hoboken) 297, 826–33 (2014).

15. Zhou, X., Marks, P.A., Rifkind, R.A. & Richon, V.M. Cloning and characterization of a histone deacetylase, HDAC9. Proc Natl Acad Sci U S A 98, 10572–7 (2001).

16. Sugo, N. et al. Nucleocytoplasmic translocation of HDAC9 regulates gene expression and dendritic growth in developing cortical neurons. Eur J Neurosci 31, 1521–32 (2010).

17. Zhang, C.L. et al. Class II histone deacetylases act as signal-responsive repressors of cardiac hypertrophy. Cell 110, 479–88 (2002).

18. Lang, B. et al. HDAC9 is implicated in schizophrenia and expressed specifically in post-mitotic neurons but not in adult neural stem cells. Am J Stem Cells 1, 31–41 (2012).

19. Malhotra, R. et al. HDAC9 is implicated in atherosclerotic aortic calcification and affects vascular smooth muscle cell phenotype. Nat Genet 51, 1580–1587 (2019).

20. Lu, J., McKinsey, T.A., Zhang, C.L. & Olson, E.N. Regulation of skeletal myogenesis by association of the MEF2 transcription factor with class II histone deacetylases. Mol Cell 6, 233–44 (2000).

21. Morrison, B.E. & D’Mello, S.R. Polydactyly in mice lacking HDAC9/HDRP. Exp Biol Med (Maywood) 233, 980–8 (2008).

22. Wilderman, A., VanOudenhove, J., Kron, J., Noonan, J.P. & Cotney, J. High-Resolution Epigenomic Atlas of Human Embryonic Craniofacial Development. Cell Rep 23, 1581–1597 (2018).

23. Minoux, M. et al. Gene bivalency at Polycomb domains regulates cranial neural crest positional identity. Science 355 (2017).

24. Andrey, G. et al. Characterization of hundreds of regulatory landscapes in developing limbs reveals two regimes of chromatin folding. Genome Res 27, 223–233 (2017).

25. Philippakis, A.A. et al. The Matchmaker Exchange: a platform for rare disease gene discovery. Hum Mutat 36, 915–21 (2015).

26. Yoon, J.G. et al. Molecular Diagnosis of Craniosynostosis Using Targeted Next-Generation Sequencing. Neurosurgery 87, 294–302 (2020).

27. De Marco, P. et al. A de novo balanced translocation t(7;12)(p21.2;p12.3) in a patient with Saethre-Chotzen-like phenotype downregulates TWIST and an osteoclastic protein-tyrosine phosphatase, PTP-oc. Eur J Med Genet 54, e478–83 (2011).

28. Karczewski, K.J. et al. The mutational constraint spectrum quantified from variation in 141,456 humans. Nature 581, 434–443 (2020).

29. Percival, C.J. et al. The effect of automated landmark identification on morphometric analyses. J Anat 234, 917–935 (2019).

30. Yang, H., Wang, H. & Jaenisch, R. Generating genetically modified mice using CRISPR/Cas-mediated genome engineering. Nat Protoc 9, 1956–68 (2014).

31. Montague, T.G., Cruz, J.M., Gagnon, J.A., Church, G.M. & Valen, E. CHOPCHOP: a CRISPR/Cas9 and TALEN web tool for genome editing. Nucleic Acids Res 42, W401–7 (2014).

32. Attanasio, C. et al. Fine tuning of craniofacial morphology by distant-acting enhancers. Science 342, 1241006 (2013).

33. Li, Q. et al. A systematic approach to identify functional motifs within vertebrate developmental enhancers. Dev Biol 337, 484–95 (2010).

34. Fisher, S. et al. Evaluating the biological relevance of putative enhancers using Tol2 transposon-mediated transgenesis in zebrafish. Nat Protoc 1, 1297–305 (2006).

35. Katz, D.C. et al. Facial shape and allometry quantitative trait locus intervals in the Diversity Outbred mouse are enriched for known skeletal and facial development genes. PLoS One 15, e0233377 (2020).

36. Percival, C.J., Marangoni, P., Tapaltsyan, V., Klein, O. & Hallgrimsson, B. The Interaction of Genetic Background and Mutational Effects in Regulation of Mouse Craniofacial Shape. G3-Genes Genomes Genetics 7, 1439–1450 (2017).

37. Adams, D.C., M. L. Collyer, A. Kaliontzopoulou, and E.K. Baken. Geomorph: Software for geometric morphometric analyses. in R package version 4.0. (2021).

38. Klingenberg, C.P. Analyzing fluctuating asymmetry with geometric morphometrics: concepts, methods, and applications. Symmetry 7, 843–934 (2015).

39. Palmer, A.R. & Strobeck, C. Fluctuating Asymmetry Analysis Unplugged. in Developmental Instability (DI): Causes and Consequences (ed. Polak, M.) 279–319 (Oxford University Press, 2003).

40. Benítez, H.A. et al. Breaking symmetry: Fluctuating asymmetry and geometric morphometrics as tools for evaluating developmental instability under diverse agroecosystems. Symmetry 12, 1789 (2020).

41. Schlager, S. Chapter 9. Morpho and Rvcg – R-packages for geometric morphometrics, shape analysis and surface manipulations. in Statistical Shape and Deformation Analysis (eds. Zheng, G., S., L. & G., S.) (Academic Press, New York, 2017).

42. Mitteroecker, P. & Gunz, P. Advances in Geometric Morphometrics. Evolutionary Biology 36, 235–247 (2009).

43. Hargrave, M., Bowles, J. & Koopman, P. In situ hybridization of whole-mount embryos. Methods Mol Biol 326, 103–13 (2006).

44. Schwartzman, O. et al. UMI-4C for quantitative and targeted chromosomal contact profiling. Nat Methods 13, 685–91 (2016).

45. Rigueur, D. & Lyons, K.M. Whole-mount skeletal staining. Methods Mol Biol 1130, 113–121 (2014).

