## Supplementary figures and tables for "*HDAC9* structural variants disrupting *TWIST1* transcriptional regulation lead to craniofacial and limb malformations"

### Supplementary data

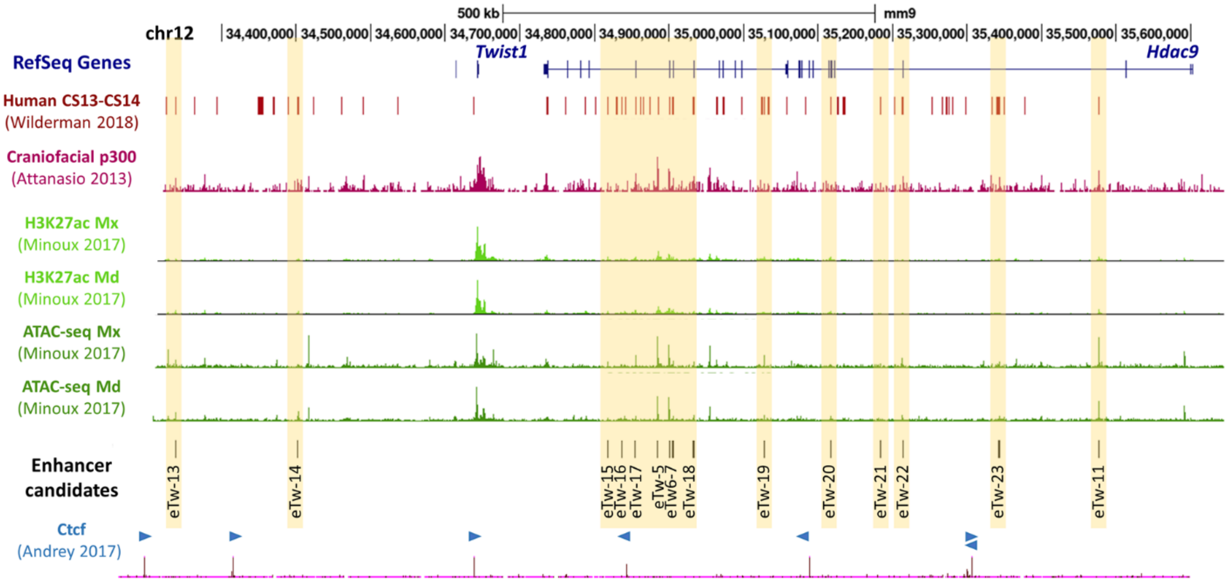


**Supplementary Figure 1**. **15 enhancer candidates at the *Twist1-Hdac9* locus.** UCSC genome browser tracks of enhancer-associated marks from early human embryonic craniofacial tissues (stages CS13-CS14) (Wilderman, 2018); p300 ChIP-seq from mouse E11.5 craniofacial tissues (Attanasio et al., 2013); H3K27ac ChIP-seq and ATAC-seq from the maxilla (Mx) and mandibula (Md) E11.5 mouse embryos (Minoux et al. 2017). Selected enhancer candidates are marked by a yellow rectangle. Ctcf ChIP-seq from mouse E11.5 limb bud (Andrey et al., 2017). Blue triangles marked bound Ctcf site and its directionality.

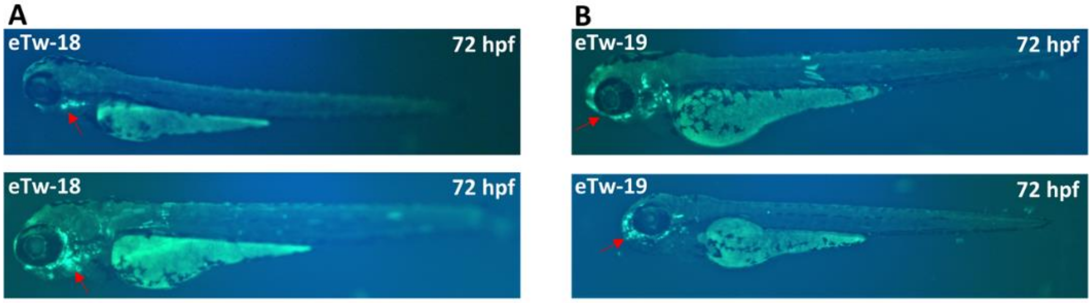


**Supplementary Figure 2**. **Functional craniofacial enhancers in the HDAC9-TWIST1 locus characterized using zebrafish. (A)** eTw-18 drives GFP expression in the pharyngeal arches (red arrow). **(B)** eTw-19 drives GFP expression in the mandibular, nasal pit, and branchial arches (red arrow). Each enhancer is represented by two different fish at 72hpf.

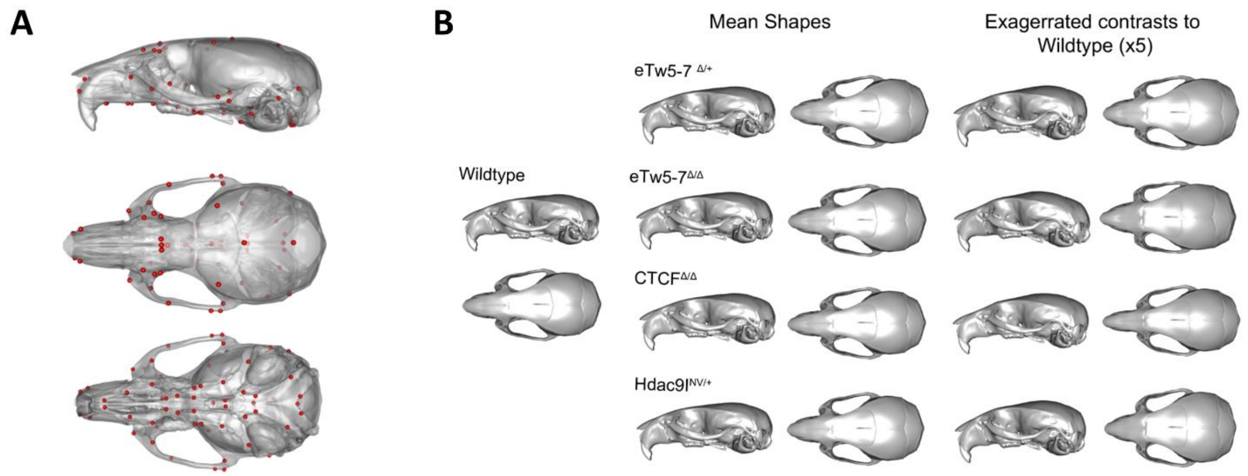


**Supplementary Figure 3**. **Skull size and shape of mouse models for *Twist1* regulatory elements (A)** 68 standardized skeletal landmarks (red dots) used to quantify 3D craniofacial form (shape and size) from micro-CT scans. **(B)** 3D morphs from micro-CT data of eTw5-7$^{\Delta/\Delta}$, eTw5-7$^{\Delta/+}$, Ctcf$^{\Delta/\Delta,}$ and *Hdac9*$^{INV/+}$ mice compared to wild-type show the differences in mean skull shape that also demonstrated by exaggerated contrasts of each model to wildtype (X5).

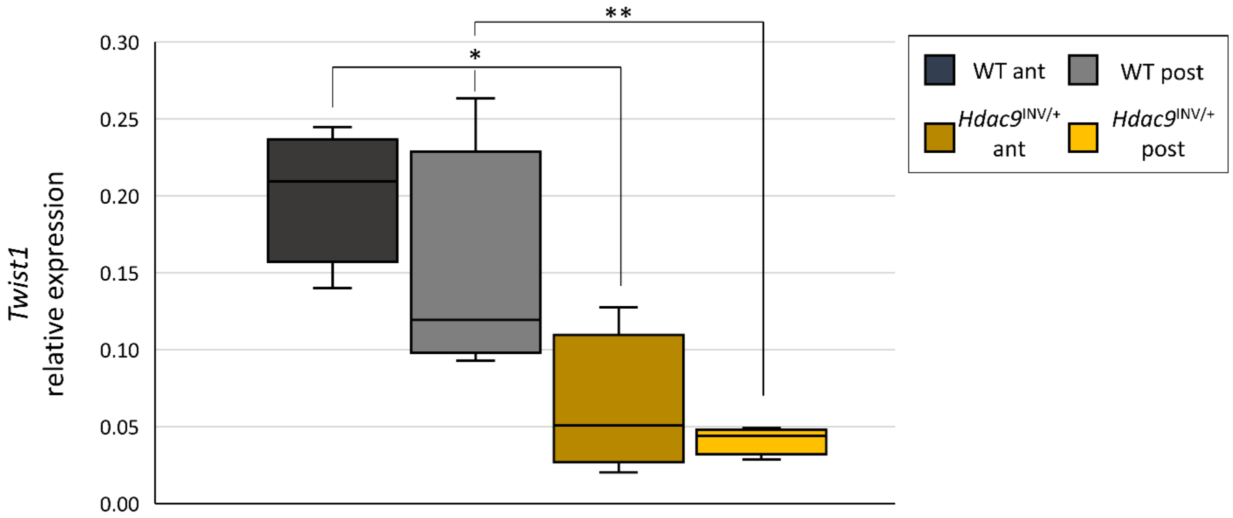


**Supplementary Figure 4:** ***Twist1*** **expression in E11.5 HL of** ***Hdac9***$^{\mathbf{INV/+}}$ **and WT mouse embryos.**

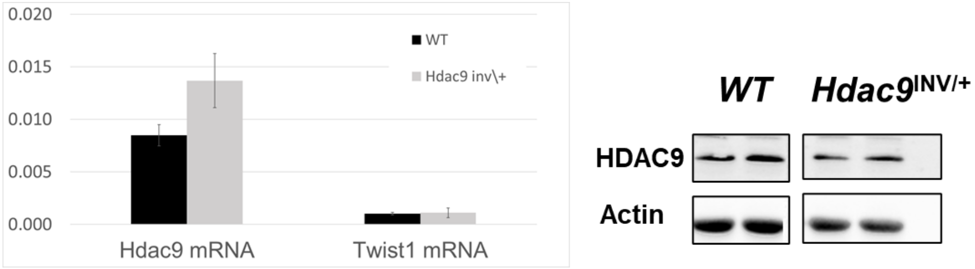


**Supplementary Figure 5: Hdac9 expression levels in adult mouse brain**. **(A)** mRNA expression levels of Hdac9 and Twist1 of adult mouse brains of wild type and *Hdac9*$^{INV/+}$**.** **(B)** Western blot analysis of Hdac9 protein level in adult mouse brains of wild type and *Hdac9*$^{INV/+}$ (Anti-Hdac9; ab59718).

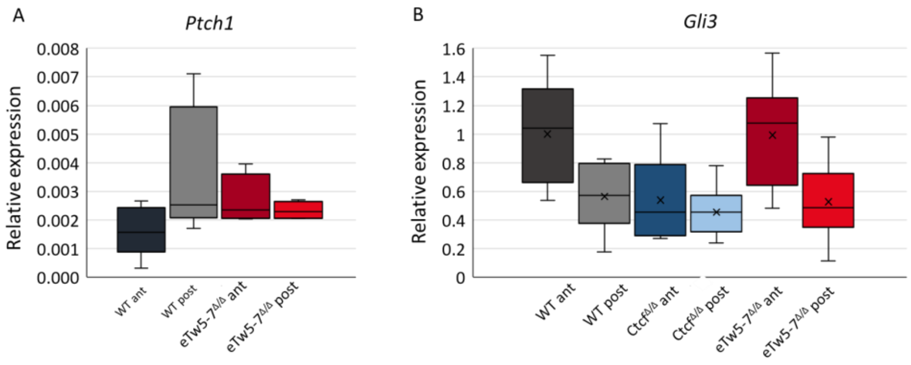


**Supplementary Figure 6: Expression pattern of Shh pathway genes**. **(A)** mRNA expression levels of *Ptch1* in HL of E11.5 WT and eTw5-7$^{\Delta/\Delta}$ mouse embryos. **(B)** mRNA expression levels of *Gli3* in HL of E11.5 WT, eTw5-7$^{\Delta/\Delta}$ and Ctcf$^{\Delta/\Delta}$ mouse embryos.

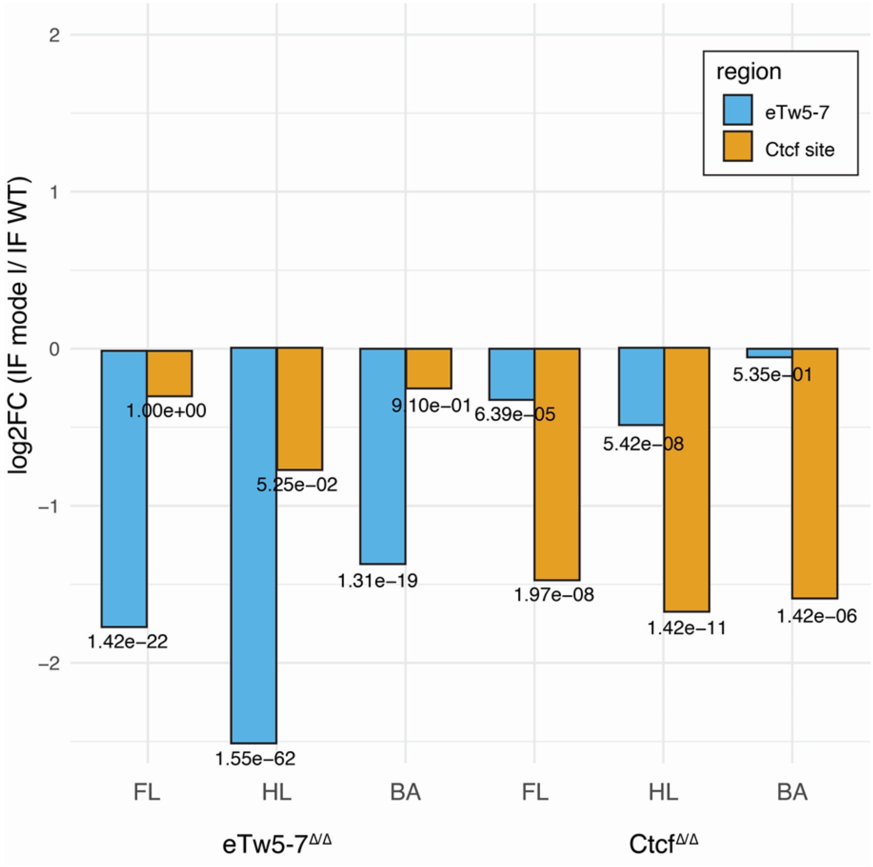


**Supplementary Figure 7: Differential interaction frequencies of eTw5-7 and Ctcf site regions with Twist1 promoter in our mouse models**. Log2 fold changes (FC) of the interaction frequency (IF) of Twist1 promoter and eTw5-7 region or Ctcf region in wildtype versus eTw5-7$^{\Delta/\Delta}$ or Ctcf $^{\Delta/\Delta}$ mice. Log2 fold changes of interaction frequencies from UMI-4C are presented for forelimb (FL), hindlimb (HL), branchial arch (BA). Statistical significance (P-value) of differential interactions was tested for these two loci (eTw5-7: chr12:34883878-34906858, Ctcf: chr12:35116684-35117991, mm9) using a Chi-square test within the UMI-4C R package (p4cIntervalsMean function).

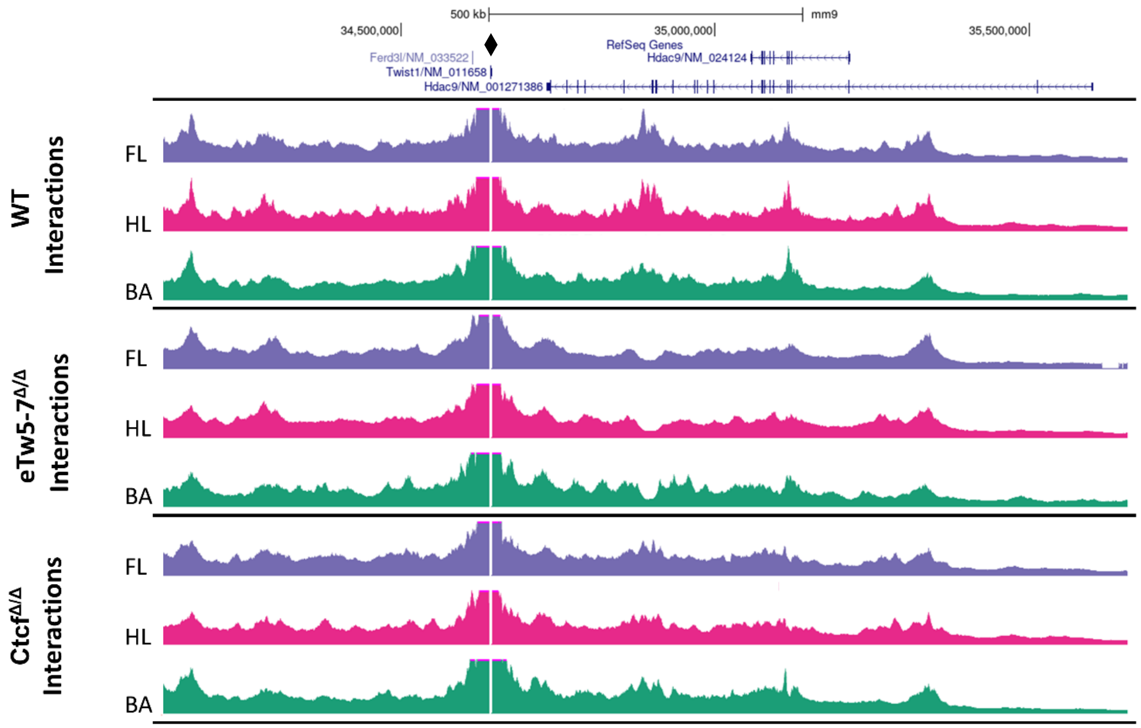


**Supplementary Figure 8: Chromatin looping in the *Hdac9-Twist1* locus in our mouse models**. UMI-4C tracks (merged from two biological replicates) of WT, eTw5-7$^{\Delta/\Delta}$ and Ctcf $^{\Delta/\Delta}$ E11.5 forelimb (FL), Hindlimb (HL), and Branchial arch (BA). The *Twist1* promoter serves as a viewpoint and is highlighted by the black diamond shape

**Supplementary Table 1:** 15 craniofacial enhancer candidates at the HDAC9-TWIST1 locus.

**Supplementary Table 2:** Zebrafish enhancer assay.

**Supplementary Table 3:** Patients with structural and nucleotide variants in the HDAC9-TWIST1 locus

**Supplementary Table 4:** MANOVA for craniofacial shape and size.

**Supplementary Table 5:** Mixed model ANOVA for fluctuating asymmetry for the pooled sample.

**Supplementary Table 6:** Pairwise absolute differences between asymmetry variances**.**

**Supplementary Table 7.** Primer list.

**Supplementary Movie 1:** Centroid size and shape of eTw5-7$^{\Delta/\Delta}$ (superior view). Here, the vector of landmark by landmark shape differences is used to quantify the difference between each genotype and the wildtype mean. Movies S1-S6 show an exaggeration (from 0-10x) of this shape difference vector to visualize the anatomical direction and distribution of shape effects by genotype.

**Supplementary Movie 2:** Centroid size and shape of eTw5-7$^{\Delta/\Delta}$ (side view).

**Supplementary Movie 3:** Centroid size and shape of Ctcf$^{\Delta/\Delta}$ (superior view).

**Supplementary Movie 4:** Centroid size and shape of Ctcf$^{\Delta/\Delta}$ (side view).

**Supplementary Movie 5:** Centroid size and shape of Hdac9$^{INV/+}$ (superior view).

**Supplementary Movie 6:** Centroid size and shape of Hdac9$^{INV/+}$ (side view).

**Table S1: 15 craniofacial enhancer candidates at the HDAC9-TWIST1 locus**

| Craniofacial TWIST1 enhacer | Human Genomic position (hg19) | mouse Genomic position (mm9) | Type (based on NM_058176) | Size (bp) |
|---|---|---|---|---|
| eTw2 | chr7:19407932-19408549 | chr12:34416711-34417304 | Intergenic | 617 |
| eTw5 | chr7:18885970-18887229 | chr12:34,884,496-34,885,742 | Intron 20 | 1260 |
| eTw6 | chr7:18874267-18875368 | chr12:34,900,126-34,901,231 | Exon 19, introns 18-19 | 1102 |
| eTw11 | chr7:18252320-18253070 | chr12:35,476,648-35,477,376 | Intergenic | 750 |
| eTw13 | chr7:19577279-19578409 | chr12:34238286-34239391 | Intergenic | 1131 |
| eTw14 | chr7:19417014-19418512 | chr12:34401724-34403227 | Intergenic | 1499 |
| eTw15 | chr7:18954896-18956568 | chr12:34817531-34819040 | Intron 21 | 1673 |
| eTw16 | chr7:18934027-18936572 | chr12:34835960-34837508 | Intron 21 | 2546 |
| eTw17 | chr7:18913392-18914585 | chr12:34854505-34855754 | Exon 21, introns 20-21 | 1194 |
| eTw18 | chr7:18832341-18833952 | chr12:34932251-34933888 | Exon 16, Introns 15-16 | 1612 |
| eTw19 | chr7:18737675-18738849 | chr12:35027281-35028407 | Intron 11 | 1175 |
| eTw20 | chr7:18629918-18630996 | chr12:35116966-35118136 | Exon 3, intron 3 | 1079 |
| eTw21 | chr7:18559965-18561240 | chr12:35183264-35184543 | Intron 1 | 1276 |
| eTw22 | chr7:18534641-18536043 | chr12:35213053-35214475 | HDAC9 5' UTR | 1403 |
| eTw23 | chr7:18391610-18393296 | chr12:35342160-35343772 | Intergenic | 1687 |

**Table S2: Zebrafish enhancer assay**

| Enhancer | Injected embryos | Positive GFP embryos at 72hpf | | | | | | | | | | | | | |
|---|---|---|---|---|---|---|---|---|---|---|---|---|---|---|---|
| | | Pectoral fin | | Branchial arch | | Caudal fin | | Brain/ Nervous system | | Heart | | Somitic muscles | | Otic vesicle | |
| *eTw_1 | 104 | 19 | 18% | 6 | 6% | 65 | 63% | 22 | 21% | 26 | 25% | 25 | 24% | - | - |
| *eTw_2 | 171 | 5 | 3% | 50 | 29% | 5 | 3% | - | - | - | - | - | - | - | - |
| *eTw_3 | 102 | - | - | - | - | 3 | 3% | 4 | 4% | - | - | 6 | 6% | - | - |
| *eTw_4 | 113 | 14 | 12% | 8 | 7% | 12 | 11% | 5 | 4% | 28 | 25% | 33 | 29% | - | - |
| *eTw_5 | 137 | 46 | 34% | 53 | 39% | 72 | 53% | 25 | 18% | 48 | 35% | 96 | 70% | - | - |
| *eTw_6 | 152 | 32 | 21% | 119 | 78% | 13 | 9% | - | - | - | - | - | - | - | - |
| *eTw_7 | 229 | 8 | 3% | - | - | - | - | - | - | - | - | - | - | - | - |
| *eTw_8 | 149 | 45 | 30% | 87 | 58% | 5 | 3% | 41 | 28% | 28 | 19% | 27 | 18% | - | - |
| *eTw_9 | 93 | 25 | 27% | - | - | 51 | 55% | - | - | - | - | - | - | - | - |
| *eTw_10 | 92 | - | - | - | - | - | - | - | - | - | - | - | - | - | - |
| *eTw_11 | 126 | 49 | 39% | 38 | 30% | 8 | 6% | 19 | 15% | 15 | 12% | 21 | 17% | 62 | 49% |
| *eTw_12 | 128 | - | - | - | - | - | - | - | - | - | - | - | - | - | - |
| eTw_13 | 109 | - | - | - | - | 2 | 2% | 2 | 2% | 2 | 2% | 2 | 2% | - | - |
| eTw_14 | 101 | 1 | 1% | 22 | 21% | 11 | 11% | 75 | 74% | 21 | 20% | 78 | 77% | - | - |
| eTw_15 | 107 | 2 | 2% | 13 | 12% | 1 | 1% | 4 | 4% | 2 | 2% | 1 | 1% | 4 | 4% |
| eTw_16 | 115 | 4 | 3% | 7 | 6% | 22 | 19% | 1 | 1% | 42 | 37% | 1 | 1% | 3 | 3% |
| eTw_17 | 160 | - | - | - | - | - | - | - | - | - | - | - | - | - | - |
| eTw_18 | 112 | - | - | 44 | 39% | 1 | 1% | 14 | 13% | - | - | 25 | 22% | 8 | 7% |
| eTw_19 | 91 | 1 | 1% | 62 | 68% | 14 | 15% | 2 | 2% | 2 | 2% | 7 | 8% | 1 | 1% |
| eTw_20 | 71 | 2 | 3% | 14 | 19% | 15 | 21% | 3 | 4% | 3 | 4% | 11 | 15% | - | - |
| eTw_21 | 152 | 4 | 3% | 22 | 14% | 7 | 4% | 33 | 22% | 40 | 26% | 82 | 54% | 4 | 2% |
| eTw_22 | 93 | - | - | 9 | 10% | - | - | 3 | 4% | 12 | 13% | 43 | 47% | - | - |
| eTw_23 | 105 | - | - | - | - | - | - | - | - | - | - | - | - | - | - |

*charcterized enhancer candidates from Hirsch et al., 2018.

### Table S3. Patients with structural and nucleotide variants in the HDAC9-TWIST1 locus

| **HDAC9 deletions** | chr. (hg19) | strart | end | type | size (Kb) | phenotype | PubMed ID |
|---|---|---|---|---|---|---|---|
| Patient 1 (Yoon et al. 2019) | chr7 | 17,298,947 | 19,057,664 | Deletion | 1,759 | Craniosynostosis, isolated sagittal and bilateral coronal CRS with facial characteristics of Saethre–Chotzen syndrome. | PMID: 31754721 |
| Patient 2 | chr7 | 18,478,923 | 18,815,377 | Deletion | 336 | A young old boy with craniosynostosis (bilateral premature fusion of the coronal sutures, underwent surgery at 6 months) and facial anomalies | Our case |
| Patient 3 | chr7 | 18,562,544 | 18,963,201 | Deletion | 401 | Craniosynostosis: scaphocephaly with anomalies of the anterior and posterior fossa; no ventricular dilatation; no parenchymal anomalies | Our case |
| Patient 5 (Lang et al. 2012) | chr7 | 17,674,100 | 18,634,682 | Deletion | 961 | Schizophrenia | PMID: 23671795 |
| Patient 6 (Lang et al. 2012) | chr7 | 18,374,606 | 18,498,502 | Deletion | 124 | Schizophrenia | PMID: 23671795 |
| Patient 7 (Lang et al. 2012) | chr7 | 18,595,436 | 18,707,719 | Deletion | 112 | Schizophrenia | PMID: 23671795 |
| **Translocation** | chr. | strart | end | | size | phenotype | PubMed ID |
| Patient 4 (De Marco, et al. 2011) | chr7 | 19,112,906 | 19,112,965 | Translocation | t(7;12)(p21.2;p12.3) | A female presented with brachycephaly with flat occipitus, ocular hypertelorism, broad nose with low nasal bridge, low-set ears, thin lower lip and small mandible. There were no associated features of the hands or feet such as syndactyly, brachydactyly, or broad first rays. The patients underwent to cranioplasty at 18-months-old. | PMID: 21708297 |
| **HDAC9 single nucleotide variants** | chr. | strart | end | | size | phenotype | PubMed ID |
| Patient 8 | chr7 | 18,631,238 | 18,631,289 | Splice site | c.542+1G>A | Autism, global developmental delays, failure to thrive and seizures, maternally-inherited HDAC9 splice-site variant. No cranosynostosis or limb malformation. | Our case |
| Patient 9 | chr7 | 18,687,535 | 18,687,536 | Frame shift | De novo, c.1166delA, p.Lys389 fs | A young male with learning impairment. Brain MRI: thin corpus callosum, septum pellucidum cyst. Also, some dysmprphic features: short stature, prominent forehead, triangular facies, prominent eyes, hypertelorism, flat phiitrum, thin upper lip. | Our case |
| Patient 10 | chr7 | 18,869,144 | 18,869,145 | Missense | s SNV, T\G, p.N813K | Young female, born from a consanguineous marriage, severe epilepsy of neonatal onset. No craniofaciel or limb phenotype. | Our case |

**Supplementary Table 4: MANOVA for craniofacial shape and size**

| | Df | SS | MS | Rsq | F | Z | Pr(>F) |
|---|---|---|---|---|---|---|---|
| Genotype | 4 | 0.017 | 0.004 | 0.172 | 11.369 | 8.902 | **0.001** |
| Sex | 2 | 0.003 | 0.002 | 0.033 | 4.368 | 3.233 | **0.003** |
| Centroid Size | 1 | 0.017 | 0.017 | 0.176 | 46.463 | 7.594 | **0.001** |
| Genotype * Sex | 4 | 0.001 | 0 | 0.015 | 0.997 | 0.138 | 0.446 |
| Genotype * Centroid Size | 4 | 0.003 | 0.001 | 0.03 | 1.961 | 2.711 | **0.008** |
| Sex * Centroid Size | 1 | 0 | 0 | 0.004 | 1.083 | 0.406 | 0.348 |
| Genotype * Sex * Centroid Size | 4 | 0.004 | 0.001 | 0.036 | 2.37 | 3.201 | **0.001** |
| Residuals | 41 | 0.052 | 0 | 0.534 | | | |
| Total | 61 | 0.098 | | | | | |

**Supplementary Table 5: Mixed model ANOVA for fluctuating asymmetry for the pooled sample.**

| Fluctuating Asymmetry (FA) ANOVA | | | | | | | | |
|---|---|---|---|---|---|---|---|---|
| | Df | SS | MS | Rsq | F | Z | Pr(>F) | |
| ind | 161 | 0.1962 | 0.0012 | 0.5751 | 2.46 | 15.75 | 0.001 | ** |
| side | 1 | 0.0654 | 0.0653 | 0.1915 | 132.11 | 7.17 | 0.001 | ** |
| FA (ind * side) | 161 | 0.0796 | 0.0005 | 0.2334 | | | | |
| Total | 323 | 0.3412 | | | | | | |

**Supplementary Table 6: Pairwise absolute differences between asymmetry variances**

| | eTw5-7 Δ/+ | eTw5-7Δ/Δ | CTCFΔ/Δ | Hdac9INV/+ |
|---|---|---|---|---|
| eTw5-7 Δ/+ | | | | |
| eTw5-7Δ/Δ | 0.000204 | | | |
| CTCFΔ/Δ | 0.000197 | 0.000007 | | |
| Hdac9INV/+ | 0.000066 | 0.000138 | 0.000131 | |
| Wildtype | 0.00026 | 0.000056 | 0.000063 | 0.000194 |

P-Values

| | eTw5-7 Δ/+ | eTw5-7Δ/Δ | CTCFΔ/Δ | Hdac9INV/+ |
|---|---|---|---|---|
| eTw5-7 Δ/+ | | | | |
| eTw5-7Δ/Δ | **0.04** | | | |
| CTCFΔ/Δ | 0.06 | 0.93 | | |
| Hdac9INV/+ | 0.54 | 0.1 | 0.13 | |
| Wildtype | **0.03** | 0.57 | 0.54 | **0.05** |

**Table S7. Primer list**

| Enhancers cloning primers | | |
|---|---|---|
| Name | Fwd primer | Rev primer |
| eTw-1 | GTTGAGTCGCTCAGATTGAGA | AGAGAAACAAAACCAAAAAAATG |
| eTw-2 | GCAGGGGGAAGAAAAAGAATA | AAGAGTAATAGCCAGAGCAGCA |
| eTw-3 | TCCACAAACTTGAGAGCGGG | CGGCACAAGCTTCTTGACC |
| eTw-4 | TAGGAAAGAAATTCTATTTATGG | AGTTCCTCCATCTTGAACTTT |
| eTw-5 | GACCCACCCTCAGACAAAAA | GGGATTTAATTTTGTAGAGTCTCCT |
| eTw-6 | ATAAAACCCCATGCCATCTTT | TATGCACTTGTTCTGCACTGG |
| eTw-7 | ATCCAAGCAATCCCTCTCCTA | GACTCAGGCGTGTATGGAAAA |
| eTw-8 | TATTTTGCGGTTTAGCTTTAA | ATGCTGGAAGCAGTAAGCTCT |
| eTw-9 | GCCCTCCATAAGGATGTGATT | AACAAGTGCCAATGGATGAAG |
| eTw-10 | CTTTGCTCAACCCAGTGCTT | TGTACCAGAAATTGGGAAGAAAA |
| eTw-11 | GATGTTGCAGTGAAGTTTCTTGT | GAGAAAATTTCTTCTCAAAGAA |
| eTw-12 | AGAACACAATGTGCCTGTTGC | CTCATCTAGGGAAGCCATAACC |
| eTw-13 | TTCTCCTTAGAACTGGGCTTT | CATGTTAAGAATGATTGGGTTTTG |
| eTw-14 | TTCAGAACCTCTTCTTTGTTTGC | GGGAATATTAAGCACGCAAA |
| eTw-15 | AGAGCATGAAAACCAAGTAGACA | CCCCAGCAATACTCAGGAAA |
| eTw-16 | TTTCACTGGCAGAGTGTGCT | TGCAGAAGTAGTTCAAAACACCT |
| eTw-17 | AAAGCCCAGTTGCATTCATT | GCTCTGAGCAAGGGGAATTA |
| eTw-18 | CCAGTTTGAACTGAAACAATCTG | GGTTTCACAAAACTCACAGTGC |
| eTw-19 | CAGGAGAAGACGCATGAACA | GGCACAGAGAAGGTTTGGAA |
| eTw-20 | CAAACAAGTTCCAAAAGCTCA | ATCGCTGGTCACAGCACATA |
| eTw-21 | GAAGACCACCTCTTGCAGGTA | TGGCTGCAAGTCAAACATAA |
| eTw-22 | GGATGGATGCGATAAAATTCA | AACCCTACAGACGCACCAAG |
| eTw-23 | TTGGAAGGTGTGTGTGGAAC | GTAACACAGGCCAGGCATTC |
| **UMI-4C primers** | | |
| *Twist1* upstream | CTGTGACAGCAGTAGTGGCA | |
| *Twist1* downstream | AATGATACGGCGACCACCGAGATCTACACTCTTTCCCTACACGACGCTCTTCCGATCTCTTCGACGCTCTGGGTGAT | |
| **gRNA and genotyping primers** | | |
| ΔeTw5-7 gRNA1 | GGGATGAGCATATGAATGG | |
| ΔeTw5-7 gRNA2 | CCATTCATATGCTCATCCC | |
| Hdac9 INV gRNA1 | GCTGGATGTGGATTCTAC | |

| | | |
|---|---|---|
| Hdac9 INV gRNA2 | GGAGGAAGTCATGTGACCTT | |
| Genotyoing WT | AGTGTCCTCAGCTCACTTGAAA | GGCCAAGTGATTTAAATAGCAAA |
| Genotyoing ΔeTw5-7 | AGTGTCCTCAGCTCACTTGAAA | AACTGTACGCAAGAAGATCAAGG |
| Genotyoing WT | GAGTTTGCTGGGAAGAAACA | GACACACAATGCATCCAGAA |
| Genotyoing *Hdac9* INV | GACACACAATGCATCCAGAA | TGTGTAGGAGCATGGTTGTG |
| **qPCR primers** | | |
| *Twist1* | CATGTCCGCGTCCCACTA | TTTTTAGTTATCCAGCTCCAGAGTC |
| *Hdac9* | GTCCCTGCCCAATATCACTCT | TGTCTGAGCATCTGTGTCTCG |
| *Gli3* | TGGGATCAGTCAGGCCATC | TGTCAGGTCCCAAATCCACAG |
| *Ptch1* | GACCGGGACTATCTGCACC | TCTCTGAAACTTCGCTCTCAG |
| *Alx4* | ACACTACCCTGATGTGTATGCA | CTCCCTCTTTCGCCACTTG |
| *Hand2* | CCGACACCAAACTCTCCAAG | TCTTGTCGTTGCTGCTCACT |
